# The bacterial virulence factors rhamnolipids and their (*R*)-3-hydroxyalkanoate precursors activate *Arabidopsis* innate immunity through two independent mechanisms

**DOI:** 10.1101/2020.12.18.423392

**Authors:** Romain Schellenberger, Jérôme Crouzet, Arvin Nickzad, Alexander Kutschera, Tim Gerster, Nicolas Borie, Corinna Dawid, Maude Cloutier, Sandra Villaume, Sandrine Dhondt-Cordelier, Jane Hubert, Sylvain Cordelier, Florence Mazeyrat-Gourbeyre, Christian Schmid, Marc Ongena, Jean-Hugues Renault, Arnaud Haudrechy, Thomas Hofmann, Fabienne Baillieul, Christophe Clément, Cyril Zipfel, Charles Gauthier, Eric Déziel, Stefanie Ranf, Stéphan Dorey

## Abstract

Plant innate immunity is activated upon perception of invasion pattern molecules by plant cell-surface immune receptors. Several bacteria of the genera *Pseudomonas* and *Burkholderia* produce rhamnolipids (RLs) from L-rhamnose and (*R*)-3-hydroxyalkanoate precursors (HAAs). RL and HAA secretion is required to modulate bacterial surface motility, biofilm development, and thus successful colonization of hosts. Here, we show that the lipidic secretome from the opportunistic pathogen *Pseudomonas aeruginosa* mostly comprising RLs and HAAs stimulates *Arabidopsis* immunity. We demonstrate that HAAs are sensed by the bulb-type lectin receptor kinase LIPOOLIGOSACCHARIDE-SPECIFIC REDUCED ELICITATION/S-DOMAIN-1-29 (LORE/SD1-29) that also mediates medium-chain 3-hydroxy fatty acid (mc-3-OH-FA) perception in the plant *Arabidopsis thaliana*. HAA sensing induces canonical immune signaling and local resistance to plant pathogenic *Pseudomonas* infection. By contrast, RLs trigger an atypical immune response and resistance to *Pseudomonas* infection independent of LORE. Thus, the glycosyl moieties of RLs, albeit abolishing sensing by LORE, do not impair their ability to trigger plant defense. In addition, our results show that RL-triggered immune response is affected by the sphingolipid composition of the plasma membrane. In conclusion, RLs and their precursors released by bacteria can both be perceived by plants but through distinct mechanisms.

**Significance:** Activation of plant innate immunity relies on the perception of microorganisms through self and nonself elicitors. Rhamnolipids and their precursor HAAs are exoproducts produced by beneficial and pathogenic bacteria. They are involved in bacterial surface dissemination and biofilm development. As these compounds are released in the extracellular milieu, they have the potential to be perceived by the plant immune system. Our work shows that both compounds independently activate plant immunity. We demonstrate that HAAs are perceived by the receptor protein kinase LORE. By contrast, rhamnolipids are not senses by LORE but activate a non-canonical immune response affected by the sphingolipid composition of the plant plasma membrane. Thus, plants are able to sense bacterial molecules as well as their direct precursors to trigger a distinct immune response.

## Introduction

Plant innate immunity activation relies on detection of invasion pattern (IP) molecules that are perceived by plant cells (1, 2). Non-self-recognition IPs include essential components of whole classes of microorganisms, such as flagellin, peptidoglycans, mc-3-OH-FAs from bacteria or chitin and β-glucans from fungi and oomycetes, respectively (3, 4). Apoplastic IPs are sensed by plant plasma membrane-localized receptor kinases (RKs) or receptor-like proteins (RLPs) that function as pattern recognition receptors (PRRs) (5, 6). Activation of the immune response requires the recruitment of regulatory receptor kinases and receptor-like cytoplasmic kinases (RLCKs) by PRRs (7). Early cellular immune signaling of pattern-triggered immunity (PTI) includes ion-flux changes at the plasma membrane, rise in cytosolic Ca^2+^ levels, production of extracellular reactive oxygen species (ROS) and activation of mitogen-activated protein kinases (MAPKs) and/or Ca^2+^-dependent protein kinases (3, 8–10). Biosynthesis and mobilization of plant hormones, including salicylic acid, jasmonic acid, ethylene, abscisic acid and brassinosteroids, ultimately modulate plant resistance to phytopathogens (11–14).

Rhamnolipids (RLs) are extracellular amphiphilic metabolites produced by several bacteria, especially *Pseudomonas* and *Burkholderia* species (15–17). Acting as wetting agents, RLs are essential for the social form of bacterial surface dissemination called swarming motility and for normal biofilm development (18–20). These glycolipids are produced from L-rhamnose and 3-(3-hydroxyalkanoyloxy)alkanoic acid (HAA) precursors (15, 21). HAAs are synthesized by dimerization of (*R*)-3-hydroxyalkanoyl-CoA in *Pseudomonas*, forming congeners through the RhlA enzyme (21). The opportunistic plant pathogen *Pseudomonas aeruginosa* and the phytopathogen *Pseudomonas syringae* produce extracellular HAAs (16, 22–24). In *P. syringae*, HAA synthesis is coordinately regulated with the late-stage flagellar gene encoding flagellin (22). HAA and RL production is finely tuned and modulates the behavior of swarming migrating bacterial cells by acting as self-produced negative and positive chemotactic-like stimuli (25). RLs contribute to the alteration of the bacterial outer membrane composition, by shedding flagellin from the flagella (26) and by releasing lipopolysaccharides (LPS) resulting in an increased hydrophobicity of the bacterial cell surface (27). In mammalian cells, RLs produced by *Burkholderia plantarii* exhibit endotoxin-like properties similar to LPS, leading to the production of proinflammatory cytokines in human mononuclear cells (28, 29). They also subvert the host innate immune response through manipulation of the human beta-defensin-2 expression (30). Moreover, RLs from *Burkholderia pseudomallei* induce Interferon gamma (IFN-γ)-dependent host immune response in goat (31).

In plants, RLs induce defense responses and resistance to biotrophic and necrotrophic pathogens (32, 33). They also contribute to the biocontrol activity of the plant beneficial bacterium *P. aeruginosa* PNA1 against oomycetes (17). Recently, it was reported that the bulb-type lectin receptor kinase LIPOOLIGOSACCHARIDE-SPECIFIC REDUCED ELICITATION/S-DOMAIN-1-29 (LORE/SD1-29) mediates medium-chain 3-hydroxy fatty acid (mc-3-OH-FA) sensing in *Arabidopsis thaliana* (hereafter, *Arabidopsis*) and that bacterial compounds comprising mc-3-OH-acyl building blocks including LPS and RLs do not stimulate LORE-dependent responses (34).

Here we show that the lipidic secretome produced by *P. aeruginosa* (RLsec) mostly composed of RLs and HAAs induces *Arabidopsis* immunity. HAAs are perceived through the RK LORE. We demonstrate that, albeit not being sensed by LORE, RLs trigger an immune response characterized by an atypical defense signature. Altogether, our results demonstrate that RLs and their precursors produced by *Pseudomonas* bacteria stimulate the plant immune response by two distinct mechanisms.

## Results

### RLsec from *Pseudomonas* induces *Arabidopsis* immune responses partially mediated by LORE

*Pseudomonas* species including opportunistic plant pathogenic or plant beneficial endophytic strains release a mixture of RL congeners and HAA precursors, here collectively termed RL secretome (RLsec) (15, 25). HPLC-MS/MS analyses of this RLsec from *P. aeruginosa* revealed the presence of mono-RLs and di-RLs at 50.9% and 44.9% of dry weight, respectively, and HAAs (3.8% of dry weight) (Supplementary Table 1). RLs comprising ten-carbon long lipid tails, Rha-C_10_-C_10_ (α-L-rhamnopyranosyl-β-hydroxydecanoyl-β-hydroxydecanoate) and Rha-Rha-C_10_-C_10_ (α-L-rhamnopyranosyl-α-L-rhamnopyranosyl-β-hydroxydecanoyl-β-hydroxydecanoate), and C_10_-C_10_ [(*R*)-3-(((*R*)-3-hydroxydecanoyl)oxy)decanoate] HAAs were the most abundant molecules in this lipidic secretome (37.6%, 33.1%, 2.1%, respectively). Notably, low amounts of free mc-3-OH-FAs (0.4% total), such as 3-OH-C_8_, 3-OH-C_10_ and 3-OH-C_12_, were also identified (Supplementary Table 1).

First, we monitored apoplastic ROS production triggered by RLsec in *Arabidopsis* (35). Wild type (WT) plants challenged with RLsec displayed a transient extracellular ROS production, starting at six minutes and peaking at 15 minutes post elicitation (Fig. 1A). A robust ROS response was detected at concentrations of RLsec starting from 0.5 μg/mL (Fig. 1B, Supplementary Fig. 1). The ROS burst was dependent on the transmembrane-NADPH oxidase RBOHD (36, 37) (Fig. 1C, Supplementary Fig. 2). RKs and RLPs mediate perception of IPs and early activation of PTI signaling (7). We monitored RLsec-triggered ROS production in *Arabidopsis* plants carrying loss-of-function mutations in genes encoding well characterized RKs and RLPs *fls2/efr1* (38, 39), *bak1*-5, *bkk1-1, bak1-5/bkk1-1* (40), *bik1/pbl1* (41), *cerk1-2* (42), *sobir1-12, sobir1-13* (43), *dorn1-1* (44) and *lore-5* (45). RLsec-induced production of ROS was only reduced in *lore-5* (Fig. 1C, Supplementary Fig. 2). Some IPs, including LPS extracts and synthetic mc-3-OH-FAs, were reported to induce a late ROS production in *Arabidopsis* (34, 46, 47). The late ROS response triggered by mc-3-OH-FAs was dependent on LORE (34). RLsec also induced a late and long-lasting ROS burst in *Arabidopsis* culminating at 6-8 hours post treatment (Fig. 2A), which was abolished in *rbohD* but not in *lore-5* mutant plants (Fig. 2A).

**Fig. 1.**
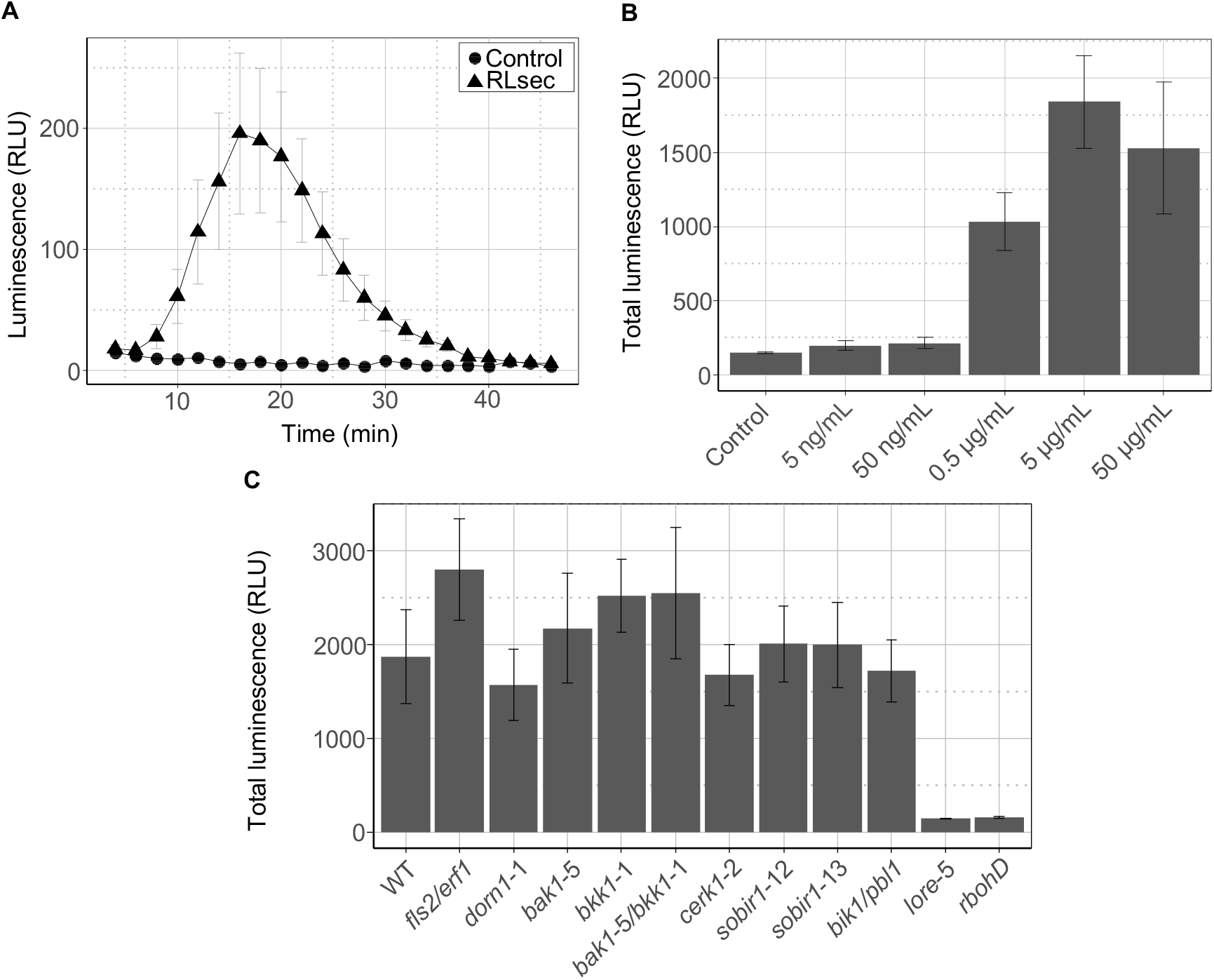
RLsec activates LORE-dependent early immune-related responses in *Arabidopsis*. (A) Extracellular ROS production after treatment of WT leaf petioles with 50 μg/mL RLsec or EtOH as control. (B) Dose effect of RLsec on ROS production. ROS production measured after treatment of WT leaf petioles with the indicated concentrations of RLsec or EtOH as control. (C) ROS production measured after treatment of WT, *fls2/efr1, dorn1-1, bak1-5, bkk1-1, bak1-5/bkk1-1, cerk1-2, sobir1-12, sobir1-13, bik1/pbl1, lore-5*, or *rbohD* leaf petioles with 50 μg/mL RLsec. **(a,b,c)** Data are mean ± SEM (n = 6). Experiments have been realized three times with similar results.

**Fig. 2.**
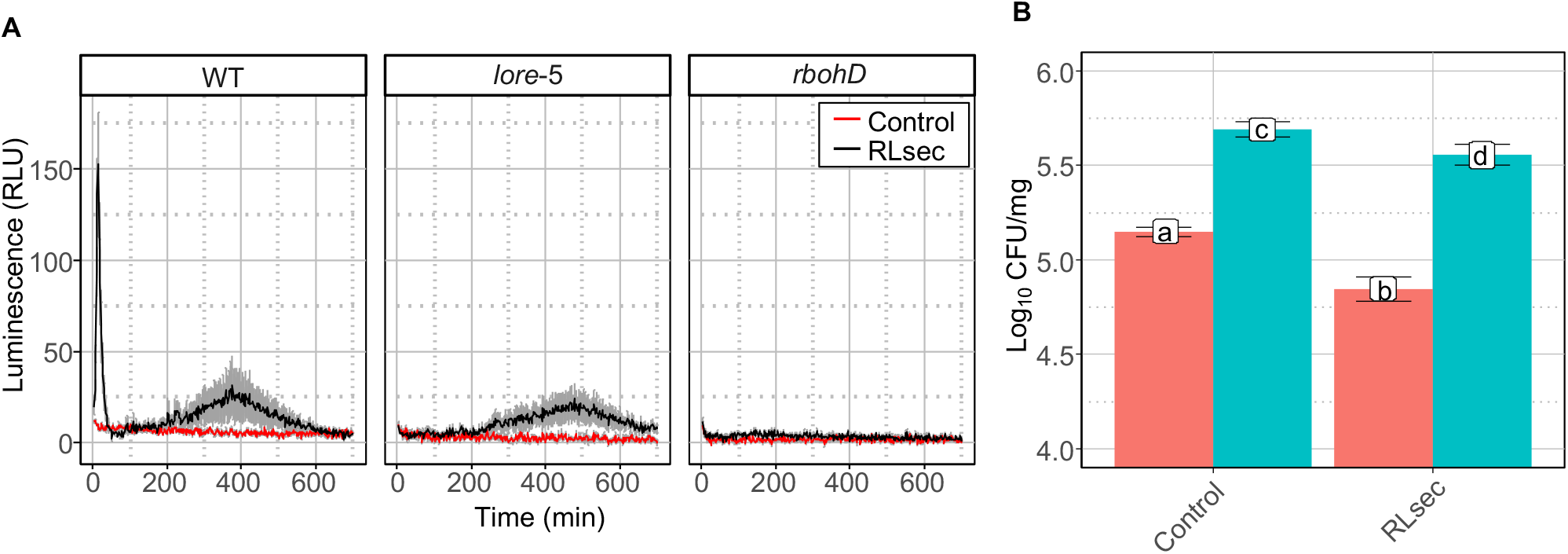
RLsec activates LORE-independent responses in *Arabidopsis*. (A) Extracellular ROS production after treatment of WT, *lore-5*, and *rbohD* leaf petioles with 50 μg/mL RLsec or EtOH (Control). ROS production was monitored over 720 minutes. Data are mean ± SEM (n = 6). Experiments have been realized three times with similar results. (B) WT (red) and *lore-5* (blue) *Arabidopsis* leaves were treated with 50 µg/mL RLsec or EtOH (control) 48 h before infection. *Pst* titers were measured at 3 d.p.i. Data are mean ± SD (n = 6, 5, 6, 6 (left to right)). Experiments have been realized twice with similar results. Letters represent results of pairwise Wilcoxon-Mann-Whitney statistic test with *P* > 0.05 (same letters) or *P* ≤ 0.05 (different letters).

Next, we tested whether RLsec induces local resistance to the hemibiotrophic phytopathogen *Pseudomonas syringae* pv. *tomato* DC3000 (*Pst*) in *Arabidopsis* (48). RLsec pretreatment significantly enhanced resistance against *Pst* infection in WT leaves and, although less pronounced, in *lore-5* plants (Fig. 2B). Taken together, our results show that RLsec induces immunity-related signaling events and disease resistance in *Arabidopsis* that are partially mediated by the bulb-type lectin RK LORE.

### *Pseudomonas* HAAs and mc-3-OH-FAs from RLsec trigger LORE-dependent *Arabidopsis* immunity

By contrast to RLsec, purified RLs do not trigger LORE-dependent [Ca^2+^]_cyt_ and early ROS signaling responses (34). Because RLsec contains significant amounts of HAAs, we investigated the role of these poorly studied compounds in RLsec-triggered immunity. We compared the responses to HAA with those to mc-3-OH-FAs, known to be sensed by LORE (34) and present in low amounts in RLsec (Supplementary table 1). Side-by-side experiments with C_10_-C_10_ HAA purified from *Pseudomonas aeruginosa* secretome and 3-OH-C_10_ revealed that both compounds induce [Ca^2+^]_cyt_ signaling and ROS production in WT plants in a dose-dependent manner (Fig. 3A and 3B, Supplementary Fig. 3 and 4). As observed upon 3-OH-C_10_ elicitation, purified C_10_-C_10_-induced ROS response was impaired in *rbohD* and *lore-5* mutants (Fig. 3C). Similarly, [Ca^2+^]_cyt_ signaling triggered by C_10_-C_10_ was impaired in *lore-5* (Fig. 3D). In addition, C_10_-C_10_ and 3-OH-C_10_ both triggered LORE-dependent MPK3 and MPK6 phosphorylation (Supplemental Fig. 5A). C_10_-C_10_ activated a late and long-lasting ROS production which, unlike the RL-triggered ROS burst, was LORE-dependent (Supplemental Fig. 6). WT but not *lore-5* mutant plants pretreated with C_10_-C_10_ or 3-OH-C_10_ displayed enhanced resistance against *Pst* (Fig. 3E). Similar to 3-OH-FAs (34), the acyl chain length of HAA affects its immune eliciting activity, as purified C_14_-_C14_ from *B. glumae* did not induce ROS production in *Arabidopsis* plants (Supplementary Fig. 7).

**Fig. 3.**
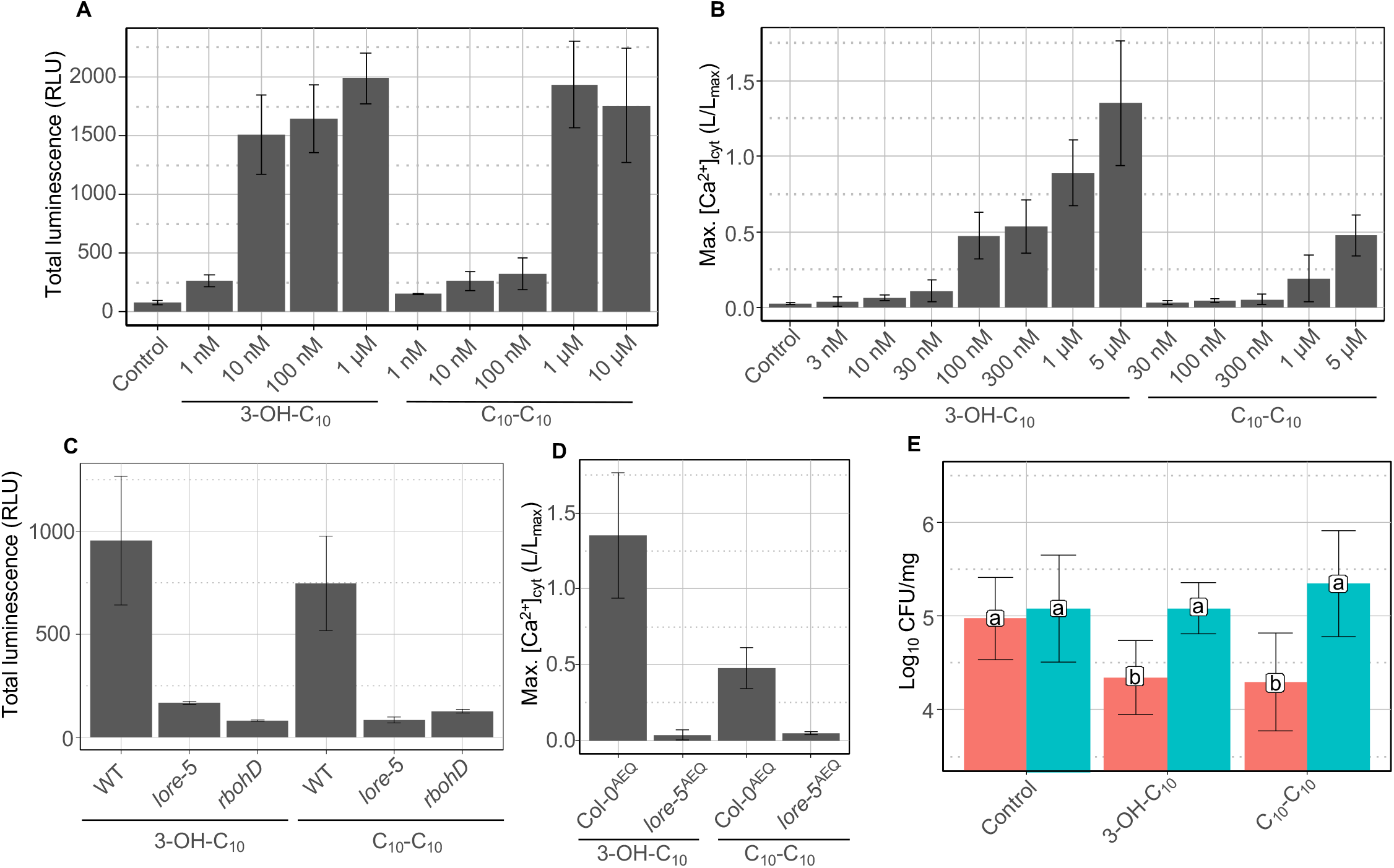
Purified HAAs from *P. aeruginosa* trigger a LORE-dependent immune response in *Arabidopsis*. (A) Dose effect of 3-OH-C_10_ and C_10_-C_10_ purified from *P. aeruginosa* on ROS production by WT leaf petioles. EtOH was used as negative control. Data are mean ± SEM (n = 6). Experiments have been realized twice with similar results. (B) Maximum (Max.) increases in [Ca^2+^]_cyt_ in *Arabidopsis* Col-0^AEQ^ seedlings treated with different concentrations of 3-OH-C_10_, C_10_-C_10_ purified from *P. aeruginosa* or MeOH as control. Data are mean ± SD (n = 3). Experiments have been realized twice with similar results. (C) ROS production measured after treatment of WT, *lore-5*, or *rbohD* leaf petioles with 10 μM 3-OH-C_10_, 10 μM purified C_10_-C_10_ or EtOH as control. Data are mean ± SEM (n = 6). Experiments have been realized three times with similar results. (D) Maximum (Max.) increases in [Ca^2+^]_cyt_ in *Arabidopsis* Col-0^AEQ^ and *lore*-5^AEQ^ seedlings treated with 5 µM 3-OH-C_10_ or purified C_10_-C_10_. Data are mean ± SD (n = 3). Experiments have been realized twice with similar results. For B and D, the same Col-0^AEQ^ 5µM data are presented (same experiments). (E) WT (red) and *lore-5* (blue) *Arabidopsis* leaves were treated with 10 μM 3-OH-C_10_, 10 μM purified C_10_-C_10_ or EtOH (control) 48 h before infection. *Pst* titers were measured at 3 d.p.i. Data are mean ± SD (n = 27, 31, 38, 13, 30, 37 (left to right)). Experiments have been realized twice with similar results. Letters represent data of pairwise Wilcoxon-Mann-Whitney statistic test with *P* > 0.05 (same letters) or *P* < 0.05 (different letters).

Trace amount of 3-OH-C_10_ was detected in C_10_-C_10_ purified from *P. aeruginosa* RLsec (Supplementary Table 2). To avoid any influence of potential contamination of HAAs with eliciting compounds related to purification procedure, we tested chemically synthesized C_10_-C_10_ for the ROS and [Ca^2+^]_cyt_ responses. Synthetic C_10_-C_10_ triggered LORE-dependent [Ca^2+^]_cyt_ signaling and ROS production in a dose-dependent manner (Fig. 4A-C). WT plants pretreated with synthetic C_10_-C_10_ also displayed LORE-dependent enhanced resistance against *Pst* infection (Fig. 4D).

**Fig. 4.**
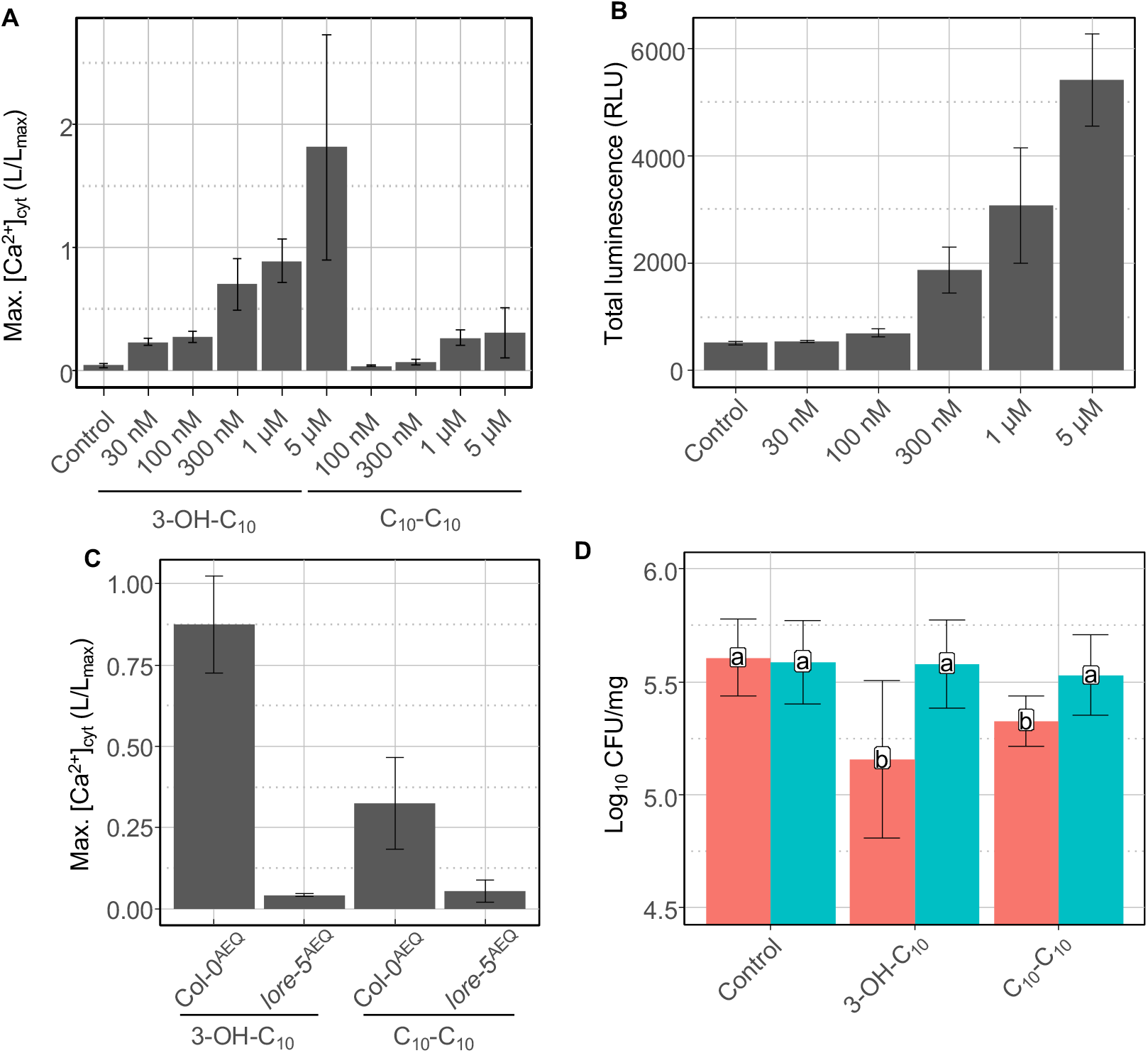
Synthetic HAAs trigger a LORE-dependent immune response in *Arabidopsis*. (A) Maximum (Max.) increases in [Ca^2+^]_cyt_ in *Arabidopsis* Col-0^AEQ^ seedlings treated with different concentrations of 3-OH-C_10_, synthetic C_10_-C_10_ or MeOH. Data are mean ± SD (n = 3). Experiments have been realized twice with similar results. (B) Dose effect of synthetic C_10_-C_10_ on ROS production by WT leaf petioles. EtOH was used as negative control. Data are mean ± SEM (n = 6). Experiments have been realized twice with similar results. (C) Maximum (Max.) increases in [Ca^2+^]_cyt_ in *Arabidopsis* Col-0^AEQ^ and *lore-5^AEQ^* seedlings treated with 5 µM 3-OH-C_10_, synthetic C_10_-C_10_ or MeOH. Data are mean ± SD (n = 3). Experiments have been realized twice with similar results. (D) WT (red) and *lore-5* (blue) *Arabidopsis* leaves were treated with 10 μM 3-OH-C_10_, 10 μM synthetic C_10_-C_10_, or MeOH (control) 48 h before infection. *Pst* titers were measured at 3 d.p.i. Data are mean ± SD (n = 17, 21, 21, 30, 14, 30 (left to right)). Experiments have been realized twice with similar results. Letters represent data of pairwise Wilcoxon-Mann-Whitney statistic test with *P* > 0.05 (same letters) or *P* ≤ 0.05 (different letters).

Altogether, our results show that HAAs secreted by *Pseudomonas* are sensed by *Arabidopsis* through the bulb-type lectin RK LORE, activate canonical PTI-related immune responses and provide resistance to bacterial infection.

### RLs trigger LORE-independent *Arabidopsis* immune responses and resistance to *Pst*

To investigate whether RLs activate a LORE-independent immune response, we used purified Rha-Rha-C_10_-C_10_ and Rha-C_10_-C_10_, the most abundant molecules from *P. aeruginosa* RLsec. In *Arabidopsis* WT, both RL congeners induced a late and long-lasting ROS production, but as observed previously (34), no early burst (Fig. 5A). As both RL congeners gave a similar ROS signature, we only used Rha-Rha-C_10_-C_10_ in the following experiments. The minimal concentration necessary to stimulate ROS production was 50 μM with an optimum at 100 μM (Fig. 5B). Late ROS production was compromised in *rbohD* but not in *lore-5* mutants (Fig. 5C). Surprisingly, neither MPK3 nor MPK6 activation by Rha-Rha-C_10_-C_10_ was detectable over a 3-hour time-course (Supplementary Fig. 5B). L-Rhamnose alone was inactive demonstrating that the lipid part of the RLs is necessary to trigger the immune response (Fig. 5A). *Burkholderia* species produce RL congeners with longer lipid chains than those produced by *Pseudomonas* (15). The RLsec from phytopathogenic *Burkholderia glumae* only contains congeners with fatty acid chain lengths varying from 12 to 16 carbons, in particular Rha-Rha-C_14_-C_14_ (49, 50). Challenge of *Arabidopsis* with purified Rha-Rha-C_14_-C_14_ from *B. glumae* did not trigger any ROS production (Fig. 5A) suggesting that the length of the fatty acid chain of RLs is critical for their eliciting activity. To determine whether RLs trigger local resistance to pathogenic *Pseudomonas* independent of LORE, plants were pretreated with 10 μM purified Rha-Rha-C_10_-C_10_ before *Pst* inoculation. WT plants displayed a significant enhanced resistance against *Pst* that was not compromised in *lore-5* mutants (Fig. 5D).

**Fig. 5.**
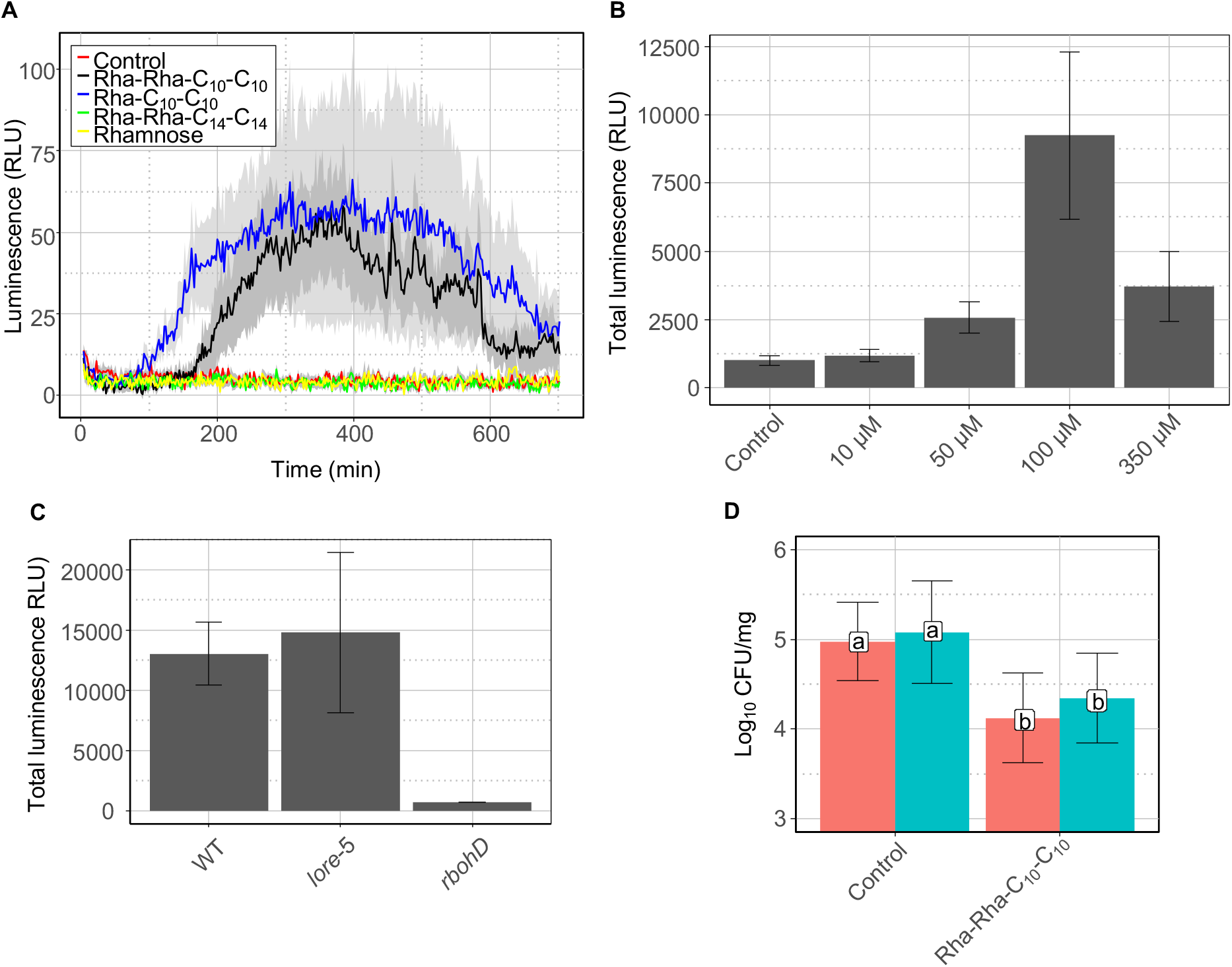
Purified RLs trigger a LORE-independent *Arabidopsis* immune response. (A) Extracellular ROS production after treatment of WT leaf petioles with 100 µM RLs, 100 µM L-rhamnose, or EtOH (control). Data are mean ± SEM (n = 6). (B) Dose effect of Rha-Rha-C_10_-C_10_ on ROS production. ROS production measured after treatment of WT leaf petioles with the indicated concentrations of Rha-Rha-C_10_-C_10_ or EtOH (control). Data are mean ± SEM (n = 6). (C) ROS production measured after treatment of WT, *lore-5*, or *rbohD* leaf petioles with 100 µM Rha-Rha-C_10_-C_10_. Data are mean ± SEM (n = 6). (D) WT (red) and *lore-5* (blue) *Arabidopsis* leaves were treated with 10 μM Rha-Rha-C_10_-C_10_ or EtOH (control) 48 h before infection. *Pst* titers were measured at 3 d.p.i. Data are mean ± SD (n = 27, 31, 30, 26 (left to right)). Letters represent data of pairwise Wilcoxon-Mann-Whitney statistic test with *P* ≤ 0.05 (same letters) or *P* < 0.05 (different letters). (A-D) Experiments have been realized three times with similar results.

To get deeper insights into the mechanisms involved in RL sensing, we used *Arabidopsis* plants carrying loss-of-function mutations in genes encoding RK and RLPs but also plasma membrane channel mutants including quintuple mechano-sensitive channels of small conductance-like (*msl4/5/6/9/10*) and double mid1-complementing activity (*mca1/2*) channel mutants (51) that could monitor changes in membrane mechanical properties. None of these mutants were affected in the long-term ROS response (Fig. 6A). Glycosylinositol phosphorylceramide (GIPC) sphingolipids were recently involved in the sensing of microbial necrosis and ethylene-inducing peptide 1-like (NLP) proteins (52). We found that the fatty acid hydroxylase *fah1/2* mutant that is disturbed in its complex sphingolipid composition (52) showed a reduced long-term ROS response (Fig. 6B). Ion leakage measurement confirmed that *fah1/2* mutant plants were less affected than WT plants by RL treatment (Fig. 6C). Ceramide synthase *loh1* mutants are also impaired in GIPC levels but not in glucosyl ceramides (52). Interestingly, RL-triggered ROS production and ion leakage was unaltered in *loh1* plants. Altogether, our results show that RLs activate an atypical immune response in *Arabidopsis* that is LORE-independent, but which is affected by the sphingolipid composition of the plasma membrane.

**Fig. 6.**
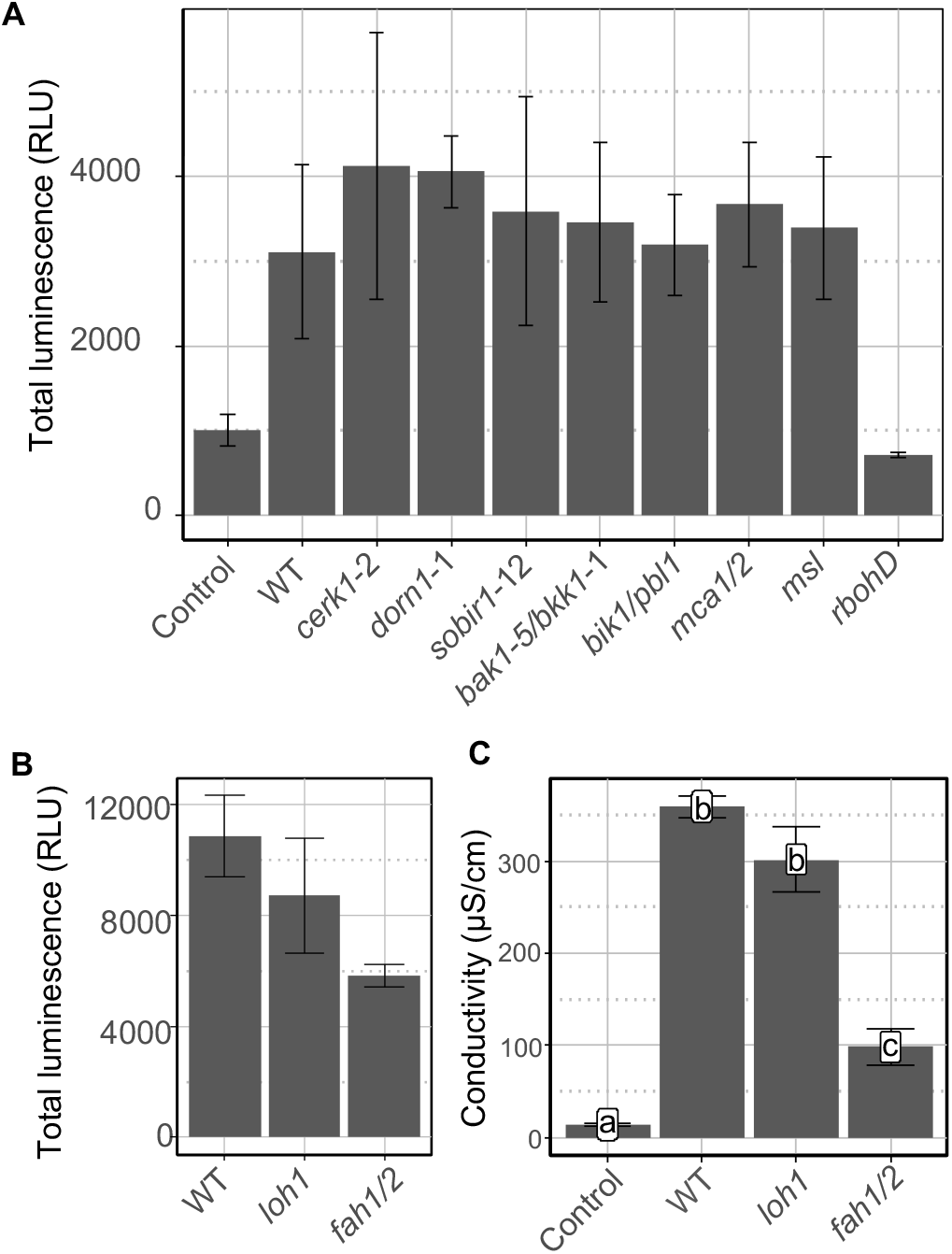
RL perception is impacted by plasma membrane sphingolipid composition. Extracellular ROS production after treatment of (A) WT, *cerk1-2, dorn1-1, sobir1-12, bak1-5/bkk1-1, bik1/pbl1, mca1/2, msl4/5/6/9/10* (*msl*), or *rbohD*, and (B) WT, *loh1*, or *fah1/2 Arabidopsis* leaf petioles with 100 µM Rha-Rha-C_10_-C_10_ or EtOH (control). Data are mean ± SEM (n = 6). Experiments have been realized three times with similar results. (C) Electrolyte leakage induced by 100 µM Rha-Rha-C_10_-C_10_ or EtOH (Control) on WT, *loh1*, or *fah1/2 Arabidopsis* leaf discs 24h post treatment. Data are mean ± SEM (n = 6). Letters represent data of pairwise Wilcoxon-Mann-Whitney statistic test with *P* ≤ 0.05 (same letters) or *P* ≤ 0.05 (different letters). Experiments have been realized twice with similar results.

## Discussion

In *Pseudomonas* and *Burkholderia* species, swarming motility is intimately related to the production of extracellular surface-active RLs and HAAs (22, 25, 53–55). In addition, RL production affects bacterial biofilm architecture and increases affinity of cells for initial adherence to surfaces through increasing the cell’s surface hydrophobicity (19, 56). These exoproducts are therefore at the frontline during host colonization. Our work demonstrates that both RLs and HAAs from the *Pseudomonas* lipidic secretome, referred to as RLsec here, are able to trigger *Arabidopsis* innate immunity by two distinct mechanisms.

We found that *Pseudomonas* RLs induce an atypical immune response. This response does not involve the RK LORE. Other bacterial compounds comprising mc-3-OH-acyl building blocks, but with large decorations including lipid A or LPS, lipopeptides, and *N*-acyl homoserine lactones also do not trigger LORE-dependent immune responses (34). RLs are glycolipids made of L-rhamnose linked to an HAA lipid tail (15, 21). Therefore, glycosylation of HAAs abolishes their perception by LORE. Glycosylation is known to affect the perception of IPs. Glycosylation of flg22 from *Acidovorax avenae* on Ser^178^ and Ser^183^ prevents its perception by rice cell (57). Similarly, unglycosylated flagellin from *Pseudomonas syringae* pv. *tabaci* 6605 induces stronger defense responses in tobacco plants than glycosylated flagellin (58). In humans, glycosylation of *Burkholderia cenocepacia* flagellin significantly reduces its perception by epithelial cells (59).

We found that RL perception does not involve previously characterized RKs, RLPs or mechanosensitive channels. However, the RL response is affected by alterations in sphingolipid synthesis suggesting a role of these key membrane lipids in RL-triggered immunity. Recently GIPCs, major structural components of the plant plasma membrane together with glucosylceramides (GlcCers), have been involved as receptors of cytotoxic NLPs (52). NLPs bind terminal monomeric hexose moieties of GIPCs. Only eudicot plants are sensing these NLPs through sphingolipid receptors. Insensitivity of monocots to NLPs is due to the length of the GIPC headgroup, consisting of three terminal hexoses compared to two in eudicots (52). *fah1/2* mutants display reduced glycosylsphingolipids (GIPCs and GlcCers) content but also lower level of ordered plasma membranes (52), suggesting that, similar to the NLP response, complex sphingolipids and/or ordered plasma membranes are necessary for the RL response. Unlike NLPs, RL responses were not significantly affected in *loh1* mutant plants also suggesting that GlcCers more than GIPC could influence RL sensing (52). Surfactin and more recently synthetic RL bolaforms and synthetic glycolipids, also active in the micromolar range, have been proposed to directly interact with plasma membrane lipids (46, 60–62). Mono- and di-RLs from *Pseudomonas* interact with phospholipids in several model membranes (63–66). In particular, RLs are able to fit into phospholipid bilayers of plant membrane model (67). In this model, the rhamnose polar heads from RLs are located near the phosphate groups from phospholipids and RL hydrophobic lipid tails are surrounded by the lipid chains from these phospholipids (67). The results obtained with these plant plasma membrane models suggest that the insertion of RLs into the lipid bilayer does not significantly affect lipid dynamics. The nature of the phytosterols could however influence the RL effect on plant plasma membrane destabilization. Subtle changes in lipid dynamics could then be linked to plant defense induction (67). Interestingly, RL bolaforms, like natural RLs are inducing a non-canonical defense signature with a long-lasting oxidative burst without MPK3 or MPK6 activation (46). This atypical defense signature triggered by two structurally different RLs, displaying amphiphilic properties and biological activities at the micromolar range, could suggest a direct interaction of these molecules with plant plasma membrane lipids.

We also demonstrated that HAAs, found in large amount in *Pseudomonas* lipidic secretome, are IPs perceived by *Arabidopsis*. HAA sensing is mediated by LORE (34). HAAs, in the micromolar range, induce typical PTI responses including transient ROS production, [Ca^2+^]_cyt_ signaling, and MPK3 and MPK6 phosphorylation in *Arabidopsis*. Interestingly, 3-OH-C_10_ activates similar responses but at concentrations 10 to 50 times lower. This is intriguing, because HAAs are present in much larger quantities (more than 3%) compared to 3-OH-FAs (0.3%) in the lipid secretome (Supplemental table 1). This high amount of HAAs could therefore compensate for their lower activity. RLs are activating an immune response at relatively high concentrations compared to both compounds. Interestingly, the RL concentration in the *P. aeruginosa* lipidic secretome is 10 to 100 times higher than HAAs and usually in the millimolar range (23, 68). RLs are produced between 20 and 110 μM *in vivo* in mammals infected by *P. aeruginosa*, especially during cystic fibrosis disease (69–71). The high concentrations of RLs needed for plant elicitation are in the range of the concentrations produced by the bacteria.

Higher steric hindrance of HAA compared to 3-OH-FAs likely results in a lower affinity to the LORE receptor. Synthetic ethyl 3-hydroxydecanoate (Et-3-OH-C10:0) and *n-*butyl 3-hydroxydecanoate (*n*But-3-OH-C10:0), which possess unbranched ester-bound carbon chains in place of the carboxyl group, also triggered LORE-dependent immune signaling in *Arabidopsis*, while 3-branched *tert*-butyl 3-hydroxydecanoate (*t*But-3-OH-C10:0) was inactive (34). HAAs, possessing a 2-branched ester-bound headgroup, activate LORE signaling. The differences in efficacy could be explained by the different steric hindrance of the molecules. Alternatively, the additional carboxyl group could account for the LORE-eliciting activity of HAAs.

*Pantoea, Dickeya* and *Pseudomonas* bacteria, in particular the well-known phytopathogen *P. syringae* mainly produce HAAs containing 3-hydroxydecanoic acid (C_10_) tails (15, 22, 72). By contrast, *Burkholderia* species including the phytopathogenic bacterium *B. glumae*, mainly produce HAAs comprising 3-hydroxytetradecanoic acid (C_14_) tails (49). *Pseudomonas* C_10_-containing HAAs activated *Arabidopsis* PTI whereas *Burkholderia* HAAs containing C_14_ fatty acid did not. Chain-length specificity was also observed for mc-3-OH-FA sensing by the LORE receptor with 3-OH-C_10_ representing the strongest immune elicitor (34). Thus, it could be hypothesized that *Arabidopsis*, and more generally *Brassicaceae* (73), are able to specifically recognize HAAs from specific bacterial species, among which several are plant opportunistic and phytopathogens (74–77). Interestingly, transcript profiles of the bean pathogen *P. syringae* pv. *syringae* B728a support a model in which leaf surface or epiphytic sites specifically favor swarming motility based on HAA surfactant production (55, 78). Low levels of HAAs contributing to motility are produced by these bacteria (22). HAA concentrations necessary to stimulate *Arabidopsis* innate immunity are in line with the concentration detected in RLsec and are produced by *Pseudomonas* (between 3 to 20% of the secretome) (23, 68, 79).

Low amounts of free mc-3-OH-FAs were found in RLsec from *P. aeruginosa* (Supplementary table 1). In *Pseudomonas*, the outer membrane lipase PagL releases 3-OH-C_10_ during synthesis of penta-acylated lipid A (34). The further fate of this 3-OH-C_10_ is unknown. RLs are able to extract LPS from the outer membrane of *P. aeruginosa* (27). Conceivably, surface-active RLs, and presumably also HAAs, could release free 3-OH-C_10_, produced through PagL activity, along with LPS from the bacterial cell wall or outer membrane vesicles (27). Alternatively, degradation of HAAs/RLs *in planta* may also release free 3-OH-C_10_. Acyl carrier protein (ACP)- and coenzyme A (CoA)-bound mc-3-OH-FAs are precursors of HAA/RL synthesis (21). Upon bacterial cell lysis, enzymatic or non-enzymatic degradation processes may also generate free 3-OH-C_10_ from these precursors. *In vivo*, insights into IP release have been recently obtained for flagellin. The plant glycosidase BGAL1 facilitates the release of immunogenic peptides from glycosylated flagellin, upstream of cleavage by proteases (80). The pathogen may evade detection by altering flagellin glycosylation and inhibiting the plant glycosidase. Flagellin glycosylation increases its physical stability that could contribute to the non-liberation/recognition of the flg22 epitope (58, 81). RLs are able to shed flagellin from *P. aeruginosa* flagella (26), suggesting that these biosurfactants participate in the release of this and presumably other eliciting compounds.

In conclusion, we hypothesize that when HAA- and RL-producing *Pseudomonas* colonize the leaf or root surface, they release RLs and HAAs which are necessary for surface motility, biofilm development, and thus successful colonization. Whereas *Arabidopsis* senses HAAs and mc-3-OH-FAs through the bulb-type lectin receptor kinase LORE, RLs are perceived through a LORE-independent mechanism. In addition to direct activation of a non-canonical defense response in plants, RLs, by releasing other IPs from bacteria, could orchestrate a node leading to strong activation of plant immunity.

## Methods

### Molecules

The *P. aeruginosa* lipidic secretome used in this study was obtained from Jeneil Biosurfactant Co., Saukville, USA (JBR-599, lot. #050629). Rha-Rha-C_10_-C_10_ and Rha-C_10_-C_10_ were purified from this lipidic secretome mixture, as previously described (33, 34). Rha-Rha-C_14_-C_14_ were purified from the *B. glumae* lipidic secretome (49). To obtain pure HAAs from *P. aeruginosa* or *B. glumae*, RLs were hydrolyzed using 1 M HCl in 1:1 dioxane-water boiling at reflux for 60 min. The mixture was extracted with ethyl acetate and the extracts were dried over anhydrous Na2SO4. After filtration, the resulting extracts were then evaporated to dryness and resuspended in 2 mL of methanol. HAAs were then isolated from digested mixture using flash chromatography on a Biotage (Stockholm, Sweden) Isorela One instrument with a SNAP Ultra C18 12g column (Biotage) using an acetonitrile/water gradient at 12 mL/min flow rate. The elution was started with 0% acetonitrile for 4.5 min and the acetonitrile concentration was raised to 100% over 28.2 min, followed by an isocratic elution of 100% acetonitrile for 13.3 min. The flash chromatography fraction containing the C_10_-C_10_ was further separated and purified using 0.25 mm thin-layer chromatographic (TLC) plates (SiliCycle SilicaPlate F-254) and developed with *n-*hexane-ethyl acetate-acetic acid (24:74:2). The bands were scraped from the plates and the HAAs, including C_10_-C_10_, were extracted from the silica with chloroform-methanol (5:1). 3-OH-C_10_ was purchased from Sigma-Aldrich Saint-Quentin Fallavier, France. All compounds were dissolved in ethanol or methanol as indicated to prepare stock solutions. Final aqueous compound dilutions were prepared freshly on the days of the experiment. Control solutions contained equal amounts of ethanol or methanol (0.05% for most experiments and not exceeding 0.5% for the highest concentrations tested). Chemical synthesis of C_10_-C_10_ is described in supplementary data 1 and 2.

### LC-MS analysis of HAAs

Samples were prepared by diluting stock solutions using MeOH to final concentration of 50 ppm. 16-Hydroxyhexadecanoic acid at 20 ppm was added to samples as internal standard^71^. The analyses were performed with a Quattro II triple quadrupole mass spectrometer (Micromass, Pointe-Claire, Canada) equipped with a Z-spray interface using electrospray ionization in negative mode. The capillary voltage was set at 3.5 kV and the cone voltage at 25 V. The source temperature was kept at 120°C and the desolvation gas at 150°C. The scanning mass range was from 130 to 930 Da. The instrument was interfaced to a high-performance liquid chromatograph (HPLC; Waters 2795, Mississauga, Ontario, Canada) equipped with a 100 x 4 mm i.d. Luna Omega PS C18 reversed-phase column (particle size 5 μm) using a water-acetonitrile gradient with a constant 2 mmol L^-1^ concentration of ammonium acetate (0.6 mL.min^-1^). Quantification of free 3-OH-C_10_ in purified C_10_-C_10_, Rha-Rha-C_10_-C_10_, Rha-C_10_-C_10_, Rha-Rha-C_14_-C_14_ or synthetic C_10_-C_10_ were performed as reported previously (34) and are presented in Supplementary table 2.

### Plant material and growth conditions

*Arabidopsis thaliana* ecotype Col-0 was used as WT parent for all experiments. Seeds from *fls2/efr1* (38, 39), *bak1-5, bkk1-1, bak1-5/bkk1-1* (40), *cerk1-2* (42), *bik1/pbl1* (41), *rbohD, msl4/5/6/9/10* and *mca1/2* (51) were provided by C. Zipfel. Seeds from *sobir1-12* and *sobir1-13* (43) were provided by F. Brunner (*Center for Plant Molecular Biology, University of Tübingen, Tübingen*, PlantResponse™). Seeds from *sd1-29* (*lore-5*), Col-0^AEQ^ and *lore*-5^AEQ^ were provided by S. Ranf (45). *loh1* and *fah1/2* seed (52) were provided by I. Feussner (*University of Göttingen, Germany*). *dorn1-1* seeds (44) were obtained from NASC stock (SALK_042209). All mutants are in the Col-0 background. Plants were grown on soil in growth chambers at 20°C, under 12 h light / 12 h dark regime with fluorescent light of 150 μmol m^-2^ s^-1^ and 60% relative humidity.

### Extracellular ROS production and calcium signaling

ROS assays were performed on 4- to 6-week-old *Arabidopsis* plants cultured on soil. Briefly, 5 mm long petiole sections were cut and placed in 150 μL of distilled water overnight in 96 wells plate (PerkinElmer) (46). Then, the protocol was conducted as previously described (82). Luminescence (relative light units, RLU) was measured every 2 min during 46 or 720 min with a Tecan Infinite F200 PRO (or a TECAN CM SPARK for Supplementary figure 6), Tecan France. Total ROS production was calculated by summing RLU measured between 4 to 46 or 4 to 720 minutes after treatment. Control was realized on petioles of WT or mutant plants. [Ca^2+^]_cyt_ measurements were done as previously described (34).

### MAPK phosphorylation assays

For MAPK phosphorylation assays, 3 leaf disks (9 mm diameter) were collected from 4 to 6-week-old *Arabidopsis* plants grown on soil and incubated 8 h in distilled water. Leaf disks were mock-treated or treated with different molecules. 15 min, 1 hour, and 3 hours after treatment, plant tissues were frozen in liquid nitrogen. To extract proteins, 60 mg of leaf tissues were ground in a homogenizer Potter-Elvehjem with 60 μL of extraction buffer (0.35 M Tris-HCl (pH 6.8), 30% (v/v) glycerol, 10% (v/v) SDS, 0.6 M DTT, 0.012% (w/v) bromophenol blue). Total protein extracts were denatured for 7 min at 95°C, centrifuged at 11 000g for 5 min and 30 μL of supernatant were separated by 12% SDS-PAGE. Proteins were transferred onto PVDF membranes for 10 min at 25 V using iBLOT gel transfer system (Invitrogen). After 30 min in 5% saturation solution (50 g L^-1^ milk, TBS (137 mM NaCl, 2.7 mM KCl, 25 mM Tris-HCl), Tween20 0.05% (v/v)) and 3 times 5 min in 0.5% washing solution (5 g L^-1^ milk, TBS (137 mM NaCl, 2.7 mM KCl, 25 mM Tris-HCl), Tween 20 0.05% (v/v)), membranes were incubated overnight with rabbit polyclonal primary antibodies against phospho-p44/42 MAPK (Erk1/2) (Cell Signaling, 1:2000) at 4°C. Then, membranes were washed 3 times 5 min with washing solution and incubated 1 h with anti-rabbit IgG HRP-conjugated secondary antibodies (Bio-Rad, 1:3000) at room temperature. Finally, washed membranes were developed with SuperSignal^®^ West Femto using an odyssey scanner (ODYSSEY^®^ Fc Dual-Mode Imaging System, LI-COR). To normalize protein loading, membranes were stripped 15 min with 0.25 M NaOH, blocked 30 min in 5% non-fat milk. Then, membranes were incubated at room temperature for 1 h with plant monoclonal anti-actin primary antibodies (CusAb, 1:1000) and 1 h with anti-mouse IgG HRP-conjugated secondary antibodies (Cell Signaling, 1:3000). Membranes were washed and developed as described above.

### Conductivity assay

The assay was performed as described previously (83), with few modifications. Eight leaf discs of 6-mm-diameter were incubated in distilled water overnight. One disc was transferred into 1.5 mL tube containing fresh distilled water and the corresponding elicitor concentration or ethanol for control. Conductivity measurements (three to four replicates for each treatment) were then conducted using a B-771 LaquaTwin (Horiba) conductivity meter.

### *Pseudomonas syringae* culture and disease resistance assays

*Pseudomonas syringae* pv. *tomato* strain DC3000 was grown at 28°C under stirring in King’s B (KB) liquid medium supplemented with antibiotics: 50 µg mL^-1^ of rifampicin and 50 µg mL^-1^ of kanamycin. For local protection assays, 15 seeds were sown per pot and grown for 3 to 5 weeks in soil. Plants were sprayed with molecules or ethanol as control and were placed two days in high humidity atmosphere before infections. Plants were inoculated by spraying the leaves with 3 mL of a bacterial suspension at an optical density (OD_600_) of 0.01 (0.025 % Silwet L-77, 10 mM MgCl2). Quantification (colony forming units) of *in planta* bacterial growth was performed 3 dpi. To this end, all plant leaves from the same pot were harvested, weighed, and crushed in a mortar with 10 mL of 10 mM MgCl_2_ and serial dilutions were performed. For each dilution, 10 μL were dropped on KB plate supplemented with appropriate antibiotics. Colony forming units (CFU) were counted after 2 days of incubation at 28°C. The number of bacteria per mg of plants fresh mass was obtained with the formula:

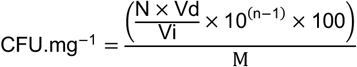

with N equal to CFU number, Vi the volume depot on plate, Vd the total volume, n the dilution number and M the plants fresh mass.

## Acknowledgments

We are thankful to Laetitia Parent and Sylvain Milot for technical support and Ralph Hückelhoven for critical discussion. This work was supported by grants from EliDeRham and Rhamnoprot (Région Grand Est). The project Rhamnoprot is co-funded by the European Union FEDER program. Work on rhamnolipids in the Déziel Lab is funded by the Natural Sciences and Engineering Research Council of Canada (NSERC) through Discovery grants RGPIN-2015-03931 and RGPIN-2020-06771. Work in the Ranf lab is supported by the German Research Foundation (SFB924/TP-B10 and Emmy Noether programme RA2541/1).

## Author contributions

J.C. and S.D. designed the research; R.S., J.C., A.K., T.G., M.T., S.V., S.D.C. performed the experiments; A.N. and E.D. purified and characterized HAAs and *B. glumae* RLs; M.C. and C.G. chemically synthetized HAAs; N.B., J.H., A.H. and J.H.R., purified *P. aeruginosa* RLs; C.D. and C.S. quantified mc-3-OH-FAs in all samples; R.S., J.C., S.C., F.M.G., F.B., S.R., E.D. and S.D. analyzed the data; R.S., J.C. and S.D. wrote the manuscript. M.O., J.H.R., A.H., T.H., C.Z., F.B., C.C., S.R and E.D contributed ideas, and critically revised the manuscript. All authors discussed the results and approved the manuscript.

## Additional Information

### Data availability

The authors declare that all data supporting the findings of this study can be found within the manuscript and its Supplementary Files. Additional data supporting the findings of this study are available from the corresponding authors upon request.

---

**Supplementary fig. 1.**
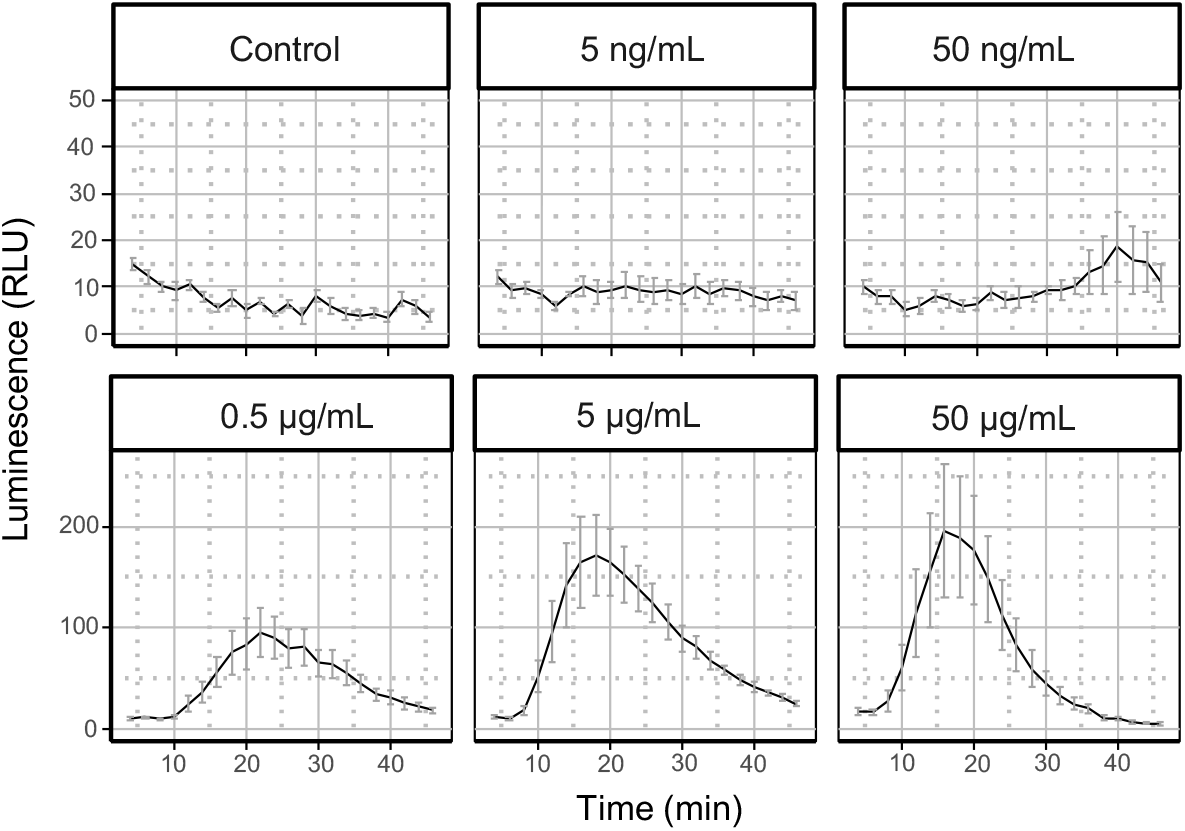
RLsec dose effect on ROS. ROS production measured after treatment of WT leaf petioles with RLsec at the indicated concentrations or EtOH (control). Data are mean ± SEM (n = 6). Experiments have been realized three times with similar results. The data presented here as kinetic are from the same experiments illustrated in fig. 1B.

**Supplementary fig. 2.**
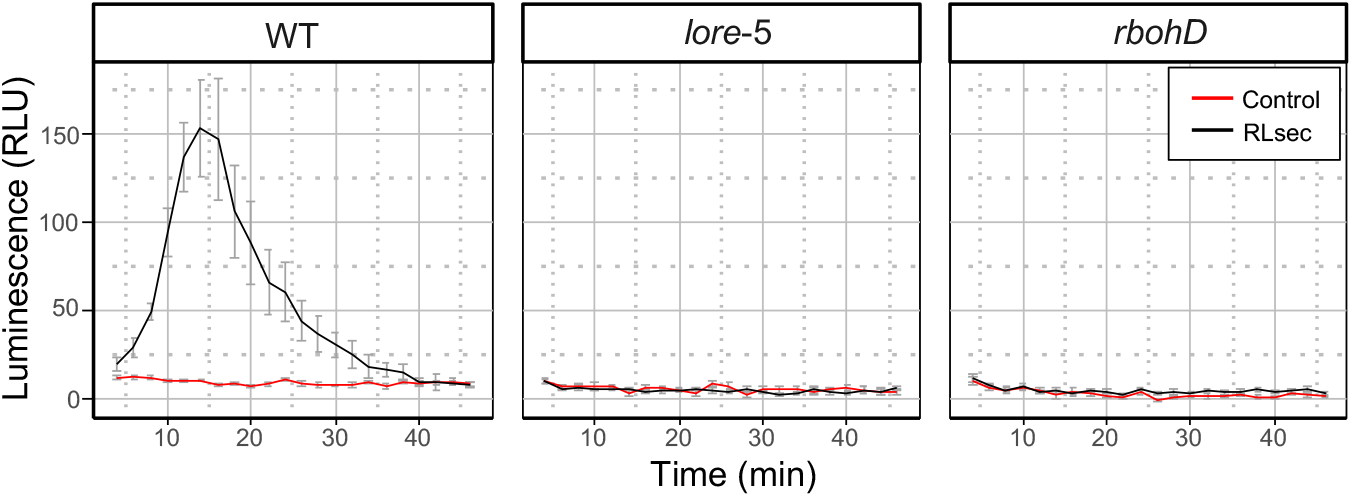
RLsec induce early ROS production through LORE and RBOHD in *Arabidopsis*. Extracellular ROS production after treatment of WT, *lore-5*, or *rbohD* leaf petioles with 50 µg/mL RLsec or EtOH (control). Data are mean ± SEM (n = 6). Experiments have been realized three times with similar results. The data presented here as kinetic are from the same experiments illustrated in fig. 1C.

**Supplementary fig. 3.**
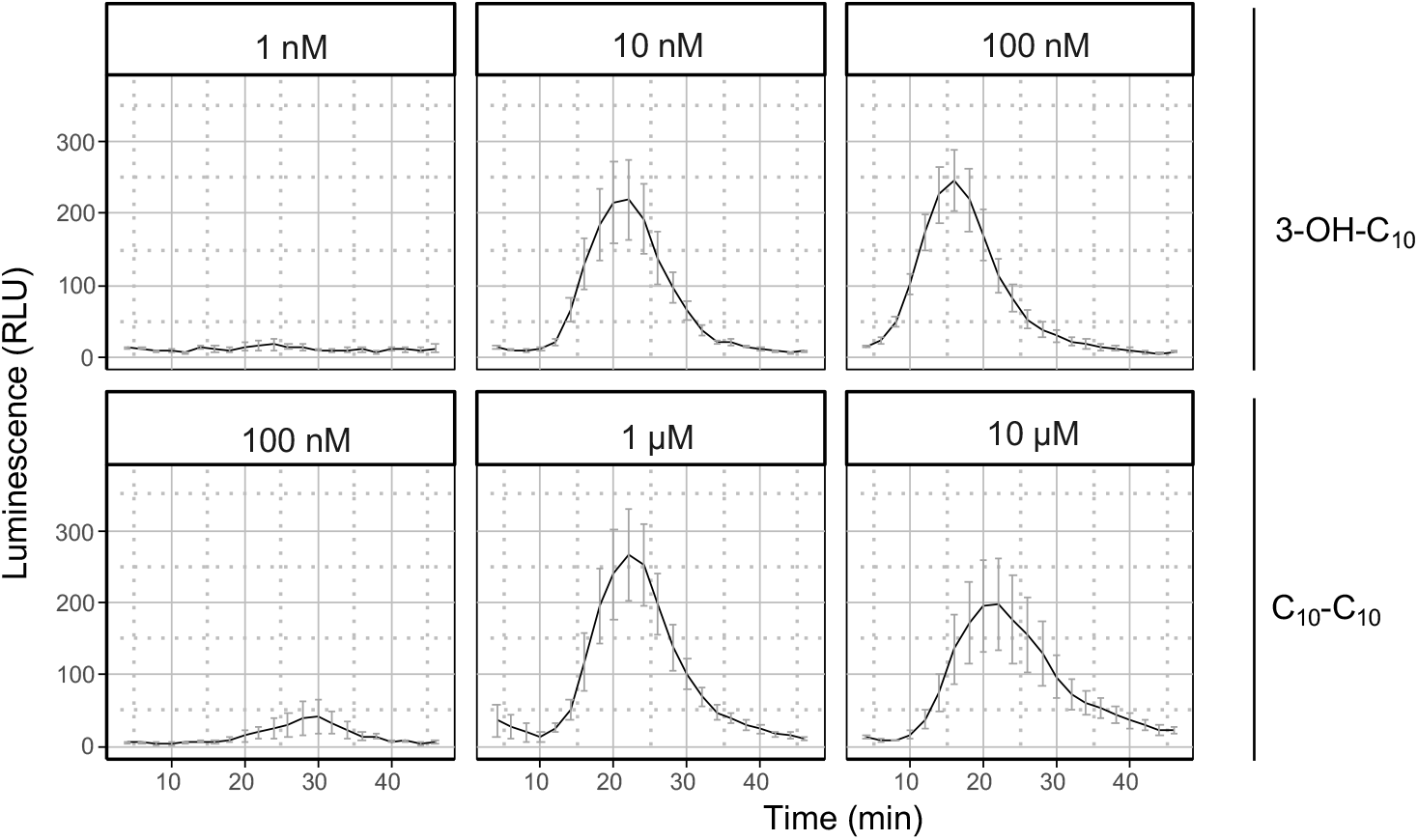
Dose effect of 3-OH-C_10_ and C_10_-C_10_ purified from *P. aeruginosa* on ROS production. ROS production measured after treatment of WT leaf petioles with the indicated concentrations of 3-OH-C_10_ and purified C_10_-C_10_. Data are mean ± SEM (n = 6). Experiments have been realized twice with similar results. The data presented here as kinetic are from the same experiments illustrated in fig. 3A.

**Supplementary fig. 4.**
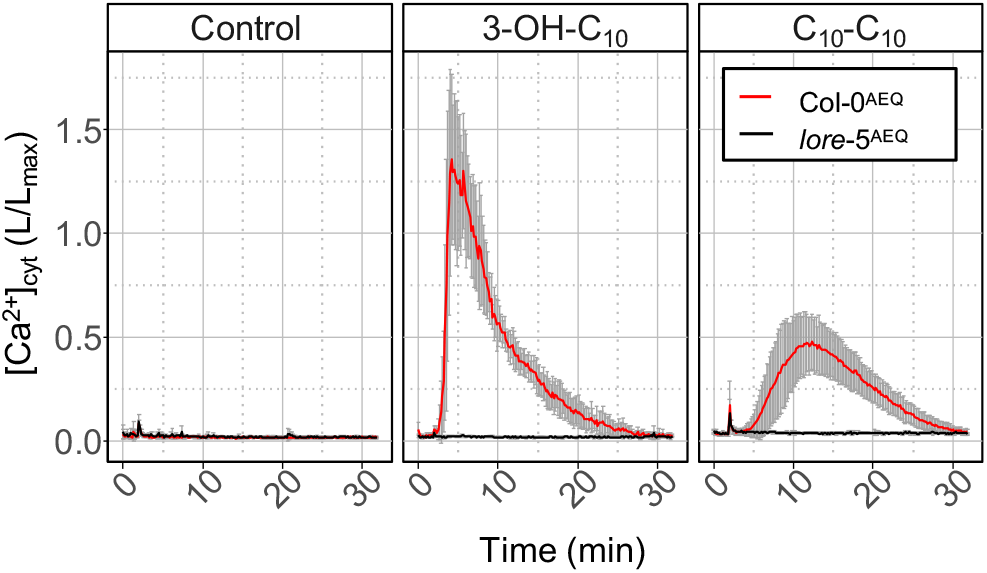
Time course Ca^2+^ signaling. [Ca^2+^]_cyt_ over time in Col-0^AEQ^ and *lore*-5^AEQ^ seedlings after treatment with 5 µM purified C_10_-C_10_, 5 µM 3-OH-C_10_ or MeOH (control). Data are mean ± SD (n = 3). Experiments have been realized twice with similar results. The data presented here as kinetic are from the same experiments illustrated in fig. 3B.

**Supplementary fig. 5.**
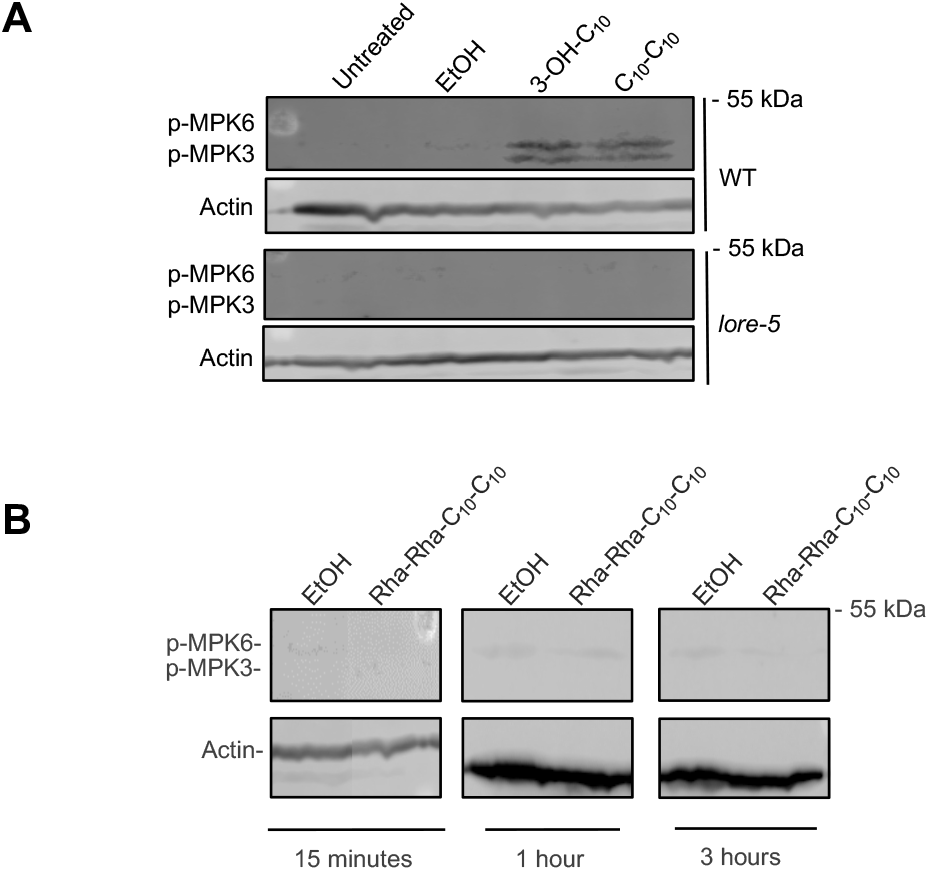
MAPK assay. Activation of MPK3 and MPK6 in (A) WT and *lore-5* leaf disk 15 minutes after treatment with 10 μM 3-OH-C_10_, 10 μM purified C_10_-C_10_ or EtOH; (B) WT leaf disk 15 minutes, 1 hour, and 3 hours after treatment with 100 µM Rha-Rha-C_10_-C_10_ or EtOH. Actin was used as loading control. Experiments have been realized twice with similar results.

**Supplementary fig. 6.**
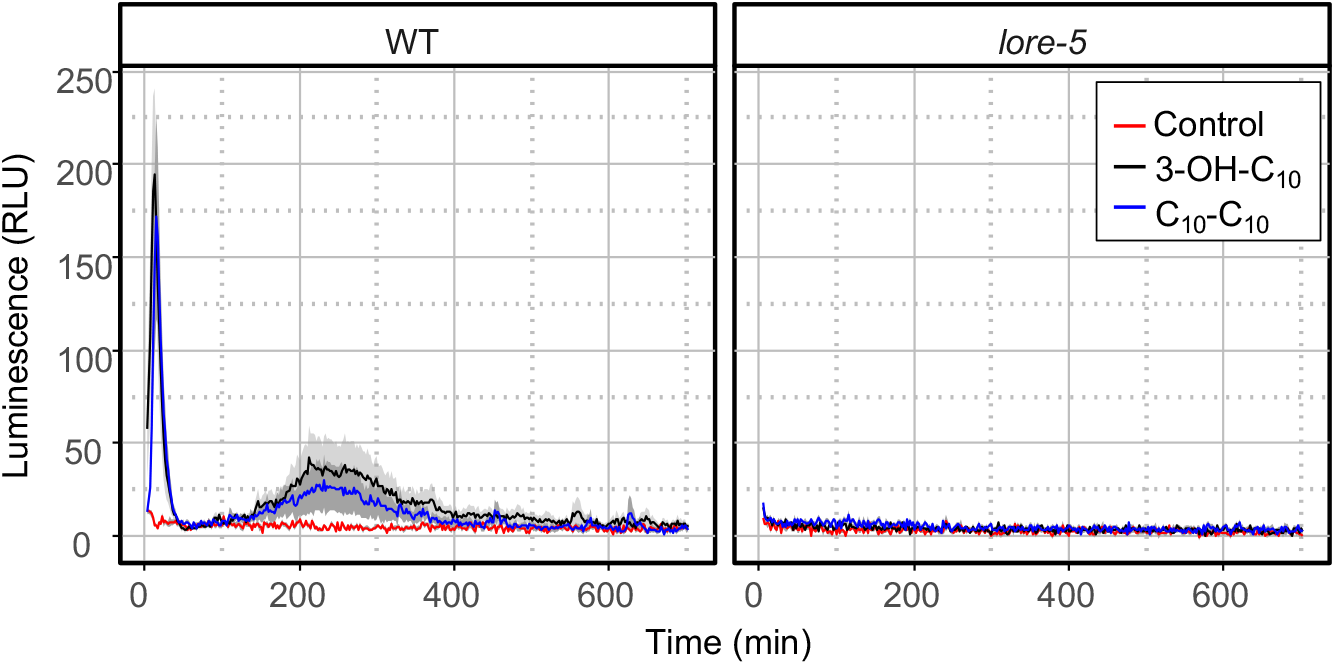
ROS production measured after treatment of WT or *lore*-5 leaf petioles with 10 μM 3-OH-C_10_, 10 μM purified C_10_-C_10_ or EtOH (control). Data are mean ± SEM (n = 6). Experiments have been realized twice with similar results.

**Supplementary fig. 7.**
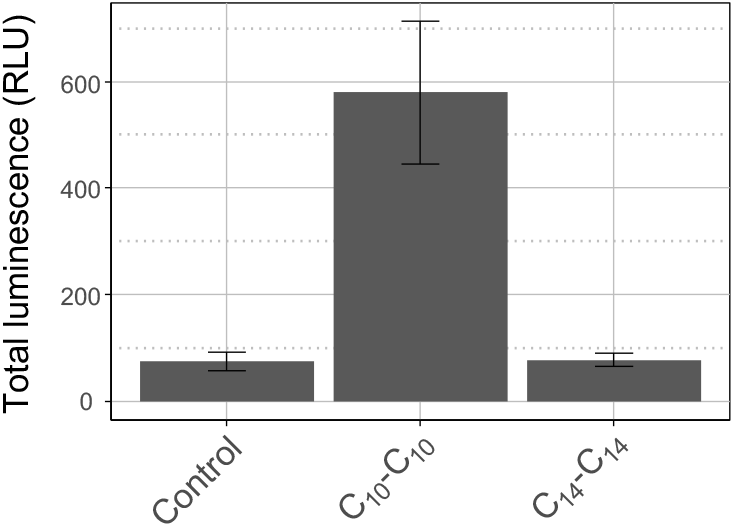
Chain length of HAAs impact *Arabidopsis* ROS immune response. ROS production measured after treatment of WT leaf petioles with 10 μM of purified C_10_-C_10_ from *Pseudomonas aeruginosa*, C_14_-C_14_ purified from *Burkholderia glumae* or with EtOH (control). Data are mean ± SEM (n = 6). Experiments have been realized three times with similar results.

**Supplementary table 1.** RLsec composition. Distribution of congeners (percent) present in the lipidic secretome produced by *P. aeruginosa* (Jeneil, JBR-599, lot. #050629).

| Molecule | % (dry weight) |
| --- | --- |
| **3-OH-C<sub>8</sub>, 3-OH-C<sub>10</sub>, 3-OH-C<sub>12</sub>** | **0.37** |
| **HAAs** | **3.75** |
| C<sub>8</sub>-C<sub>8</sub> | 0.23 |
| C<sub>8</sub>-C<sub>10</sub> | 0.85 |
| C<sub>10</sub>-C<sub>10</sub> | 2.13 |
| C<sub>10</sub>-C<sub>12</sub> | 0.26 |
| C<sub>12</sub>-C<sub>12</sub> | 0 |
| C<sub>8</sub>-C<sub>12:1</sub> | 0.09 |
| C<sub>10</sub>-C<sub>12:1</sub> | 0.12 |
| C<sub>12</sub>-C<sub>12:1</sub> | 0.07 |
| **monorhamnolipids** | **50.94** |
| Rha-C<sub>8</sub>-C<sub>8</sub> | 0 |
| Rha-C<sub>8</sub>-C<sub>10</sub> | 5.28 |
| Rha-C<sub>10</sub>-C<sub>10</sub> | 37.61 |
| Rha-C<sub>10</sub>-C<sub>12</sub> | 3.53 |
| Rha-C<sub>12</sub>-C<sub>12</sub> | 0.06 |
| Rha-C<sub>8</sub>-C<sub>12:1</sub> | 0.72 |
| Rha-C<sub>10</sub>-C<sub>12:1</sub> | 3.55 |
| Rha-C<sub>12</sub>-C<sub>12:1</sub> | 0.19 |
| **dirhamnolipids** | **44.94** |
| Rha-Rha-C<sub>8</sub>-C<sub>8</sub> | 0.36 |
| Rha-Rha-C<sub>8</sub>-C<sub>10</sub> | 4.33 |
| Rha-Rha-C<sub>10</sub>-C<sub>10</sub> | 33.14 |
| Rha-Rha-C<sub>10</sub>-C<sub>12</sub> | 4.22 |
| Rha-Rha-C<sub>12</sub>-C<sub>12</sub> | 0.10 |
| Rha-Rha-C<sub>8</sub>-C<sub>12:1</sub> | 0.64 |
| Rha-Rha-C<sub>10</sub>-C<sub>12:1</sub> | 1.88 |
| Rha-Rha-C<sub>12</sub>-C<sub>12:1</sub> | 0.27 |

**Supplementary table 2.** Quantification of free 3-OH-C_10_ in HAA and RL samples.

| Molecules | Origin | Reference | Sample concentration (for quantification) | Free 3-OH-C<sub>10</sub> concentration | Sample concentrations (for biological assays) | Concentration of 3-OH-C<sub>10</sub> at the compound concentration used for the experiments | MTI in *Arabidopsis* | LORE-dependent |
| --- | --- | --- | --- | --- | --- | --- | --- | --- |
| **Mono-RL (Rha-C<sub>10</sub>-C<sub>10</sub>)** | *Pseudomonas aeruginosa* PA14 | 36 | 5 mM | 0.28 µM | 100 µM | 5 nM | Yes | No |
| **Di-RL (Rha-Rha-C<sub>10</sub>-C<sub>10</sub>)** | *Pseudomonas aeruginosa* PA14 | 36 | 5 mM | 0.05 µM | 10 µM to 350 µM | 100 pM to 3 nM | Yes | No |
| **Di-RL (Rha-Rha-C<sub>14</sub>-C<sub>14</sub>)** | *Burkholderia glumae* | 53 | nd | nd | 100 µM | nd | No | No |
| **C<sub>14</sub>-C<sub>14</sub>** | *Burkholderia glumae* | this study | 1 mM | <LOQ | 10 µM | <LOQ | No | No |
| **C<sub>10</sub>-C<sub>10</sub>** | *Pseudomonas aeruginosa* PA14 | this study | 0.1 mM | 0.09 µM | 1 nM to 10 µM | 0.9 pM to 9 nM | Yes | Yes |
| **Synthetic C<sub>10</sub>-C<sub>10</sub>** | Chemical synthesis (see Supplementary data 1 and 2) | this study | 0.1 mM | <LOQ | 30 nM to 10 µM | <LOQ | Yes | Yes |

## References

1. D. E. Cook, C. H. Mesarich, B. P. Thomma, Understanding plant immunity as a surveillance system to detect invasion. Annu. Rev. Phytopathol. 53, 541–563 (2015).

2. K. Kanyuka, J. J. Rudd, Cell surface immune receptors: the guardians of the plant’s extracellular spaces. Curr. Opin. Plant Biol. 50, 1–8 (2019).

3. T. Boller, G. Felix, A renaissance of elicitors: perception of microbe-associated molecular patterns and danger signals by pattern-recognition receptors. Annu. Rev. Plant Biol. 60, 379–406 (2009).

4. M. A. Newman, T. Sundelin, J. T. Nielsen, G. Erbs, MAMP (microbe-associated molecular pattern) triggered immunity in plants. Front. Plant Sci. 4, 139 (2013).

5. F. Boutrot, C. Zipfel, Function, discovery, and exploitation of plant pattern recognition receptors for broad-spectrum disease resistance. Annu. Rev. Phytopathol. 55, 257–286 (2017).

6. S. Ranf, Sensing of molecular patterns through cell surface immune receptors. Curr. Opin. Plant Biol. 38, 68–77 (2017).

7. D. Couto, C. Zipfel, Regulation of pattern recognition receptor signalling in plants. Nat. Rev. Immunol. 16, 537–552 (2016).

8. J. Bigeard, J. Colcombet, H. Hirt, Signaling mechanisms in pattern-triggered immunity (PTI). Mol. Plant 8, 521–539 (2015).

9. A. Garcia-Brugger et al., Early signaling events induced by elicitors of plant defenses. Mol. Plant-Microbe Interact. 19, 711–724 (2006).

10. S. Wu, L. Shan, P. He, Microbial signature-triggered plant defense responses and early signaling mechanisms. Plant Sci. 228, 118–126 (2014).

11. D. De Vleesschauwer, G. Gheysen, M. Hofte, Hormone defense networking in rice: tales from a different world. Trends Plant Sci. 18, 555–565 (2013).

12. J. Glazebrook, Contrasting mechanisms of defense against biotrophic and necrotrophic pathogens. Annu. Rev. Phytopathol. 43, 205–227 (2005).

13. A. Robert-Seilaniantz, M. Grant, J. D. Jones, Hormone crosstalk in plant disease and defense: more than just jasmonate-salicylate antagonism. Annu. Rev. Phytopathol. 49, 317–343 (2011).

14. L. Trda et al., Perception of pathogenic or beneficial bacteria and their evasion of host immunity: pattern recognition receptors in the frontline. Front. Plant Sci. 6, 219 (2015).

15. A. M. Abdel-Mawgoud, F. Lépine, E. Déziel, Rhamnolipids: diversity of structures, microbial origins and roles. Appl. Microbiol. Biotechnol. 86, 1323–1336 (2010).

16. V. U. Irorere, L. Tripathi, R. Marchant, S. McClean, I. M. Banat, Microbial rhamnolipid production: a critical re-evaluation of published data and suggested future publication criteria. Appl. Microbiol. Biotechnol. 101, 3941–3951 (2017).

17. M. Perneel et al., Phenazines and biosurfactants interact in the biological control of soil-borne diseases caused by *Pythium* spp. Environ. Microbiol. 10, 778–788 (2008).

18. L. Chrzanowski, L. Lawniczak, K. Czaczyk, Why do microorganisms produce rhamnolipids? World J. Microbiol. Biotechnol. 28, 401–419 (2012).

19. A. Nickzad, E. Déziel, The involvement of rhamnolipids in microbial cell adhesion and biofilm development - an approach for control? Lett. Appl. Microbiol. 58, 447–453 (2014).

20. P. Vatsa, L. Sanchez, C. Clément, F. Baillieul, S. Dorey, Rhamnolipid biosurfactants as new players in animal and plant defense against microbes. Int. J. Mol. Sci. 11, 5095–5108 (2010).

21. A. M. Abdel-Mawgoud, F. Lépine, E. Déziel, A stereospecific pathway diverts beta-oxidation intermediates to the biosynthesis of rhamnolipid biosurfactants. Chem. Biol. 21, 156–164 (2014).

22. A. Y. Burch et al., Pseudomonas syringae coordinates production of a motility-enabling surfactant with flagellar assembly. J. Bacteriol. 194, 1287–1298 (2012).

23. E. Déziel, F. Lépine, S. Milot, R. Villemur, *rhlA* is required for the production of a novel biosurfactant promoting swarming motility in *Pseudomonas aeruginosa:* 3-(3-hydroxyalkanoyloxy)alkanoic acids (HAAs), the precursors of rhamnolipids. Microbiology 149, 2005–2013 (2003).

24. J. M. Plotnikova, L. G. Rahme, F. M. Ausubel, Pathogenesis of the human opportunistic pathogen *Pseudomonas aeruginosa* PA14 in *Arabidopsis*. Plant Physiol. 124, 1766–1774 (2000).

25. J. Tremblay, A. P. Richardson, F. Lépine, E. Déziel, Self-produced extracellular stimuli modulate the *Pseudomonas aeruginosa* swarming motility behaviour. Environ. Microbiol. 9, 2622–2630 (2007).

26. U. Gerstel, M. Czapp, J. Bartels, J. M. Schroder, Rhamnolipid-induced shedding of flagellin from *Pseudomonas aeruginosa* provokes hBD-2 and IL-8 response in human keratinocytes. Cell. Microbiol. 11, 842–853 (2009).

27. R. A. Al-Tahhan, T. R. Sandrin, A. A. Bodour, R. M. Maier, Rhamnolipid-induced removal of lipopolysaccharide from *Pseudomonas aeruginosa:* effect on cell surface properties and interaction with hydrophobic substrates. Appl. Environ. Microbiol. 66, 3262–3268 (2000).

28. J. Andrä et al., Endotoxin-like properties of a rhamnolipid exotoxin from *Burkholderia (Pseudomonas) plantarii:* immune cell stimulation and biophysical characterization. Biol. Chem. 387, 301–310 (2006).

29. J. Bauer, K. Brandenburg, U. Zahringer, J. Rademann, Chemical synthesis of a glycolipid library by a solid-phase strategy allows elucidation of the structural specificity of immunostimulation by rhamnolipids. Chemistry 12, 7116–7124 (2006).

30. J. Dossel, U. Meyer-Hoffert, J. M. Schroder, U. Gerstel, *Pseudomonas aeruginosa-derived* rhamnolipids subvert the host innate immune response through manipulation of the human beta-defensin-2 expression. Cell. Microbiol. 14, 1364–1375 (2012).

31. M. Gonzalez-Juarrero et al., Polar lipids of *Burkholderia pseudomallei* induce different host immune responses. PloS one 8, e80368 (2013).

32. L. Sanchez et al., Rhamnolipids elicit defense responses and induce disease resistance against biotrophic, hemibiotrophic, and necrotrophic pathogens that require different signaling pathways in *Arabidopsis* and highlight a central role for salicylic acid. Plant Physiol. 160, 1630–1641 (2012).

33. A. L. Varnier et al., Bacterial rhamnolipids are novel MAMPs conferring resistance to *Botrytis cinerea* in grapevine. Plant, Cell Environ. 32, 178–193 (2009).

34. A. Kutschera et al., Bacterial medium-chain 3-hydroxy fatty acid metabolites trigger immunity in *Arabidopsis* plants. Science 364, 178–181 (2019).

35. J. Qi, J. Wang, Z. Gong, J. M. Zhou, Apoplastic ROS signaling in plant immunity. Curr. Opin. Plant Biol. 38, 92–100 (2017).

36. Y. Kadota, K. Shirasu, C. Zipfel, Regulation of the NADPH oxidase RBOHD during plant immunity. Plant Cell Physiol. 56, 1472–1480 (2015).

37. M. A. Torres, J. L. Dangl, J. D. Jones, *Arabidopsis* gp91phox homologues AtrbohD and AtrbohF are required for accumulation of reactive oxygen intermediates in the plant defense response. Proc. Natl. Acad. Sci. USA 99, 517–522 (2002).

38. D. Chinchilla, Z. Bauer, M. Regenass, T. Boller, G. Felix, The *Arabidopsis* receptor kinase FLS2 binds flg22 and determines the specificity of flagellin perception. Plant Cell 18, 465–476 (2006).

39. C. Zipfel et al., Perception of the bacterial PAMP EF-Tu by the receptor EFR restricts *Agrobacterium-mediated* transformation. Cell 125, 749–760 (2006).

40. M. Roux et al., The *Arabidopsis* leucine-rich repeat receptor-like kinases BAK1/SERK3 and BKK1/SERK4 are required for innate immunity to hemibiotrophic and biotrophic pathogens. Plant Cell 23, 2440–2455 (2011).

41. L. Li et al., The FLS2-associated kinase BIK1 directly phosphorylates the NADPH oxidase RbohD to control plant immunity. Cell Host Microbe 15, 329–338 (2014).

42. A. Miya et al., CERK1, a LysM receptor kinase, is essential for chitin elicitor signaling in *Arabidopsis*. Proc. Natl. Acad. Sci. USA 104, 19613–19618 (2007).

43. W. Zhang et al., *Arabidopsis* receptor-like protein30 and receptor-like kinase suppressor of BIR1-1/EVERSHED mediate innate immunity to necrotrophic fungi. Plant Cell 25, 4227–4241 (2013).

44. J. Choi et al., Identification of a plant receptor for extracellular ATP. Science 343, 290–294 (2014).

45. S. Ranf et al., A lectin S-domain receptor kinase mediates lipopolysaccharide sensing in *Arabidopsis thaliana*. Nat. Immunol. 16, 426–433 (2015).

46. P. Luzuriaga-Loaiza et al., Synthetic rhamnolipid bolaforms trigger an innate immune response in *Arabidopsis thaliana*. Sci. Rep. 8, 8534 (2018).

47. K. Shang-Guan et al., Lipopolysaccharides trigger two successive bursts of reactive oxygen species at distinct cellular locations. Plant Physiol. 176, 2543–2556 (2018).

48. X. F. Xin, S. Y. He, *Pseudomonas syringae* pv. tomato DC3000: a model pathogen for probing disease susceptibility and hormone signaling in plants. Annu. Rev. Phytopathol. 51, 473–498 (2013).

49. S. G. Costa, E. Déziel, F. Lépine, Characterization of rhamnolipid production by *Burkholderia glumae*. Lett. Appl. Microbiol. 53, 620–627 (2011).

50. J. H. Ham, R. A. Melanson, M. C. Rush, *Burkholderia glumae:* next major pathogen of rice? Mol. Plant Pathol. 12, 329–339 (2011).

51. A. B. Stephan, H. H. Kunz, E. Yang, J. I. Schroeder, Rapid hyperosmotic-induced Ca^2+^ responses in *Arabidopsis thaliana* exhibit sensory potentiation and involvement of plastidial KEA transporters. Proc. Natl. Acad. Sci. USA 113, E5242–5249 (2016).

52. T. Lenarcic et al., Eudicot plant-specific sphingolipids determine host selectivity of microbial NLP cytolysins. Science 358, 1431–1434 (2017).

53. N. C. Caiazza, R. M. Shanks, G. A. O’Toole, Rhamnolipids modulate swarming motility patterns of *Pseudomonas aeruginosa*. J. Bacteriol. 187, 7351–7361 (2005).

54. A. Nickzad, F. Lépine, E. Déziel, Quorum sensing controls swarming motility of *Burkholderia glumae* through regulation of rhamnolipids. PloS one 10, e0128509 (2015).

55. X. Yu et al., Transcriptional responses of *Pseudomonas syringae* to growth in epiphytic versus apoplastic leaf sites. Proc. Natl. Acad. Sci. USA 110, E425–434 (2013).

56. M. E. Davey, N. C. Caiazza, G. A. O’Toole, Rhamnolipid surfactant production affects biofilm architecture in *Pseudomonas aeruginosa* PAO1. J. Bacteriol. 185, 1027–1036 (2003).

57. H. Hirai et al., Glycosylation regulates specific induction of rice immune responses by *Acidovorax avenae* flagellin. J. Biol. Chem. 286, 25519–25530 (2011).

58. F. Taguchi et al., Glycosylation of flagellin from *Pseudomonas syringae* pv. tabaci 6605 contributes to evasion of host tobacco plant surveillance system. Physiol. Mol. Plant Pathol. 74, 11–17 (2009).

59. A. Hanuszkiewicz et al., Identification of the flagellin glycosylation system in *Burkholderia cenocepacia* and the contribution of glycosylated flagellin to evasion of human innate immune responses. J. Biol. Chem. 289, 19231–19244 (2014).

60. G. Henry, M. Deleu, E. Jourdan, P. Thonart, M. Ongena, The bacterial lipopeptide surfactin targets the lipid fraction of the plant plasma membrane to trigger immune-related defence responses. Cell. Microbiol. 13, 1824–1837 (2011).

61. M. N. Nasir et al., Differential interaction of synthetic glycolipids with biomimetic plasma membrane lipids correlates with the plant biological response. Langmuir 33, 9979–9987 (2017).

62. M. Robineau et al., Synthetic mono-rhamnolipids display direct antifungal effects and trigger an innate immune response in tomato against *Botrytis cinerea*. Molecules 25, 3108 (2020).

63. H. Abbasi, K. A. Noghabi, A. Ortiz, Interaction of a bacterial monorhamnolipid secreted by *Pseudomonas aeruginosa* MA01 with phosphatidylcholine model membranes. Chem. Phys. Lipids 165, 745–752 (2012).

64. F. J. Aranda et al., Thermodynamics of the interaction of a dirhamnolipid biosurfactant secreted by *Pseudomonas aeruginosa* with phospholipid membranes. Langmuir 23, 2700–2705 (2007).

65. A. Ortiz, F. J. Aranda, J. A. Teruel, Interaction of dirhamnolipid biosurfactants with phospholipid membranes: a molecular level study. Adv. Exp. Med. Biol. 672, 42–53 (2010).

66. M. Sanchez, F. J. Aranda, J. A. Teruel, A. Ortiz, Interaction of a bacterial dirhamnolipid with phosphatidylcholine membranes: a biophysical study. Chem. Phys. Lipids 161, 51–55 (2009).

67. N. Monnier et al., Exploring the dual interaction of natural rhamnolipids with plant and fungal biomimetic plasma membranes through biophysical studies. Int. J. Mol. Sci. 20, 1009 (2019).

68. F. Lépine, E. Déziel, S. Milot, R. Villemur, Liquid chromatographic/mass spectrometric detection of the 3-(3-hydroxyalkanoyloxy) alkanoic acid precursors of rhamnolipids in *Pseudomonas aeruginosa* cultures. J. Mass Spectrom. 37, 41–46 (2002).

69. R. Kownatzki, B. Tummler, G. Doring, Rhamnolipid of *Pseudomonas aeruginosa* in sputum of cystic fibrosis patients. Lancet 1, 1026–1027 (1987).

70. R. C. Read et al., Effect of *Pseudomonas aeruginosa* rhamnolipids on mucociliary transport and ciliary beating. J. Appl. Physiol. 72, 2271–2277 (1992).

71. M. Somerville et al., Release of mucus glycoconjugates by *Pseudomonas aeruginosa* rhamnolipid into feline trachea *in vivo* and human bronchus *in vitro. Am. J. Respir*. Cell Mol. Biol. 6, 116–122 (1992).

72. A. Germer et al., Exploiting the natural diversity of RhlA acyltransferases for the synthesis of the rhamnolipid precursor 3-(3-Hydroxyalkanoyloxy)alkanoic acid. Appl. Environ. Microbiol. 86, e02317–02319 (2020).

73. S. Ranf, Immune sensing of lipopolysaccharide in plants and animals: same but different. PLOS Pathog. 12, e1005596 (2016).

74. S. Compant, J. Nowak, T. Coenye, C. Clément, E. Ait Barka, Diversity and occurrence of *Burkholderia* spp. in the natural environment. FEMS Microbiol. Rev. 32, 607–626 (2008).

75. E. Kay, F. Bertolla, T. M. Vogel, P. Simonet, Opportunistic colonization of *Ralstonia solanacearum-infected* plants by *Acinetobacter* sp. and its natural competence development. Microb. Ecol. 43, 291–297 (2002).

76. M. W. Silby, C. Winstanley, S. A. Godfrey, S. B. Levy, R. W. Jackson, *Pseudomonas* genomes: diverse and adaptable. FEMS Microbiol. Rev. 35, 652–680 (2011).

77. I. K. Toth, L. Pritchard, P. R. J. Birch, Comparative genomics reveals what makes an enterobacterial plant pathogen. Annu. Rev. Phytopathol. 44, 305–336 (2006).

78. X. Yu et al., Transcriptional analysis of the global regulatory networks active in *Pseudomonas syringae* during leaf colonization. mBio 5, e01683–01614 (2014).

79. K. Zhu, C. O. Rock, RhlA converts beta-hydroxyacyl-acyl carrier protein intermediates in fatty acid synthesis to the beta-hydroxydecanoyl-beta-hydroxydecanoate component of rhamnolipids in *Pseudomonas aeruginosa*. J. Bacteriol. 190, 3147–3154 (2008).

80. P. Buscaill et al., Glycosidase and glycan polymorphism control hydrolytic release of immunogenic flagellin peptides. Science 364, eaav0748 (2019).

81. F. Taguchi et al., Effects of glycosylation on swimming ability and flagellar polymorphic transformation in *Pseudomonas syringae* pv. *tabaci* 6605. J. Bacteriol. 190, 764–768 (2008).

82. J. M. Smith, A. Heese, Rapid bioassay to measure early reactive oxygen species production in *Arabidopsis* leave tissue in response to living *Pseudomonas syringae*. Plant Methods 10, 6 (2014).

83. M. Magnin-Robert et al., Modifications of sphingolipid content affect tolerance to hemibiotrophic and necrotrophic pathogens by modulating plant defense responses in *Arabidopsis*. Plant Physiol. 169, 2255–2274 (2015).

84. M. De Vleeschouwer et al., Rapid total synthesis of cyclic lipodepsipeptides as a premise to investigate their self-assembly and biological activity. Chem. Eur. J. 20, 7766–7775 (2014).

