## Supplementary data 1 for "The bacterial virulence factors rhamnolipids and their (*R*)-3-hydroxyalkanoate precursors activate *Arabidopsis* innate immunity through two independent mechanisms"

Chemical synthesis of HAA C_10_-C_10_

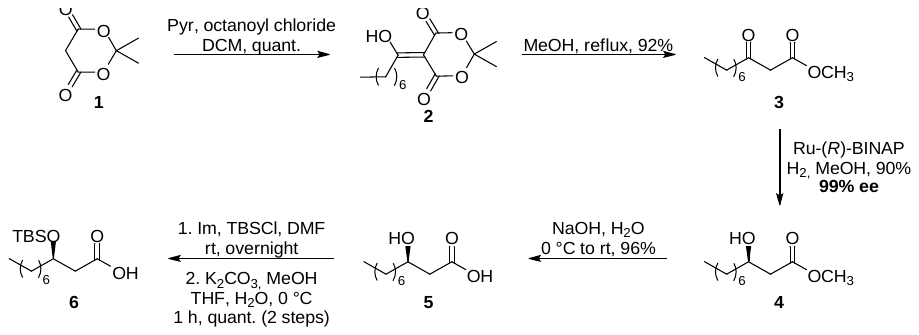

**Scheme 1.** Synthesis of acid **6** from Meldrum’s acid **1** as described by De Vleeschouwer (84).

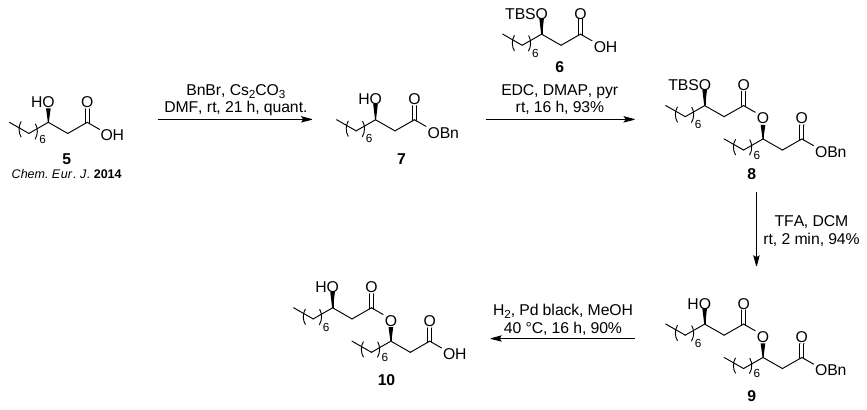

**Scheme 2.** Synthesis of C_10_-C_10_ dilipid **10** from known lipid **5** (84).

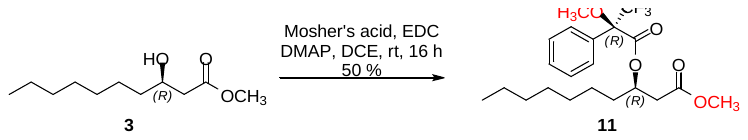

**Scheme 3.** Synthesis of Mosher’s ester **8** for the NMR determination of the enantiomeric purity (29)

All starting materials and reagents were purchased from commercial sources and used as received without further purification. Air and water sensitive reactions were performed in oven-dried glassware under Ar atmosphere. Moisture sensitive reagents were introduced via a dry syringe. Anhydrous solvents were either prepared from commercial solvents and dried over heat-gun activated 4 Å molecular sieves or supplied over molecular sieves and used as received. Reactions were monitored by thin-layer chromatography (TLC) with silica gel 60 F_254_ 0.25 mm pre-coated aluminum foil plates. Compounds were visualized by using UV_254_ and/or orcinol (1 mg mL^-1^) in 10% aq. H_2_SO_4_ solution and/or Hanessian’s stain and/or KMnO_4_ with heating. Normal-phase flash column chromatography was performed on silica gel 60 Å (15-40 μm). NMR spectra were recorded at 297 K in the indicated solvent (CDCl_3_) with a 600 MHz instrument, employing standard softwares given by the manufacturer. ^1^H and ^13^C NMR spectra were referenced to tetramethylsilane (TMS, *δ*_H_ = *δ*_C_ = 0.00 ppm) as internal reference for spectra in CDCl_3._ Assignments were based on ^1^H, ^13^C, COSY, and HSQC experiments. High-resolution mass spectra (HRMS) were recorded on an ESI-Q-TOF mass spectrometer. Optical rotations [α]^20^_D_ were measured on an Anton Paar polarimeter. The retention factor (*R_f_*) was calculated from silica gel 60 F_254_ 0.25 mm pre-coated glass TLC plates. Analytical reverse phase HPLC analysis was performed on a Thermo Fisher Scientific Ultimate 3000 HPLC-CAD system equipped with a Dionex LPG-3400SD pump, a WPS-3000SL autosampler, a TCC-3000SD column oven and a charged aerosol detector (CAD) Corona Veo. The power function value was set at 1.0, the filter at 1 s, the data collection rate at 10 Hz and the evaporated temperature at 35 °C. Nitrogen (57.2 psi) was used for nebulization. All data were analyzed using the Thermo Fischer Chromeleon 7.2.9 software. A reverse phase column Hypersil Gold (250 x 4.6 mm) was used with a mobile phase consisting of acetonitrile-water gradient containing 0.1% formic acid. Before injection, the column was equilibrated for 10 minutes with 50% of acetonitrile. The injection volume was 100 μL. The elution gradient started from 50% acetonitrile to 60% for 20 minutes, then to 100% within the next 40 minutes and hold for 15 minutes. HPLC flow rate was 0.8 mL min^-1^ and oven temperature was set at 28 °C.

**2,2-Dimethyl-5-octanoyl-1,3-dioxane-4,6-dione (2)**

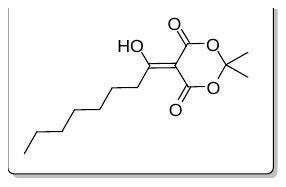

To a solution of Meldrum’s acid **1** (7443 mg, 51.64 mmol, 1.05 eq) in anhydrous DCM (22 mL) was slowly added anhydrous pyridine (8.0 mL, 98 mmol, 2.0 eq). A solution of octanoyl chloride (8.4 mL, 49 mmol, 1.0 eq) in anhydrous DCM (15 mL) was added dropwise to the reaction flask, which was then stirred at rt for 1 h under Ar atmosphere. The suspension was washed with 1 M aq. HCl (2×) and the aqueous layers were extracted with DCM (2×). The combined organic layers were washed with 1 M aq. HCl (2×) and brine, dried over anhydrous MgSO_4_, filtered, and evaporated under reduced pressure to give compound **2** (13.3 g, quant.) as a brown-red oil. The product was characterized by ^1^H NMR and the data were found to be in good agreement with those published previously (84).

**Methyl 3-oxodecanoate (3)**

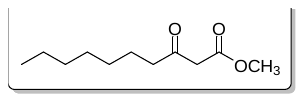

Enol **2** (12805 mg, 47.369 mmol, 1.0 eq) was solubilized in anhydrous MeOH (57 mL) and the solution was refluxed for 3 h under Ar atmosphere. The solution was then cooled at rt and evaporated under reduced pressure. The residue was purified by silica gel flash chromatography (*n-*Hex/EtOAc 100:0 to 95:0) to give keto-ester **3** (8765 mg, 92%) as a colorless oil. The product was characterized by ^1^H NMR and the data were found to be in good agreement with those published previously (84).

**(*R*)-Methyl 3-hydroxydecanoate (4)**

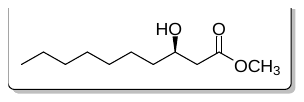

1. Preparation of the catalyst: In a heat gun dried reaction flask under Ar atmosphere were added (*R*)-BINAP (379 mg, 0.608 mmol, 0.024 eq) and (COD)Ru(2-methylallyl)_2_ (162 mg, 0.507 mmol, 0.020 eq). The reactants were then solubilized in anhydrous acetone (25 mL), which was previously degassed with Ar. A solution of 48% aq. HBr (0.13 mL) in degassed anhydrous MeOH (6.3 mL) was added to the reaction flask, and the mixture was stirred at rt for 30 min under an Ar atmosphere. The solvents were then evaporated under nitrogen flux and the catalyst was used without further purification.

2. Preparation of hydroxy-ester **4**: To the previously prepared catalyst was cannulated a solution of keto-ester **3** (5072 mg, 25.33 mmol, 1.00 eq) in anhydrous degassed MeOH (51 mL). The solution was stirred at 55 °C for 18 h under H_2_ atmosphere. The solution was then cooled to 0 °C, filtered over Celite, and evaporated under reduced pressure. The residue was purified by silica gel flash chromatography (*n-*Hex/EtOAc 9:1 to 8:2) to give hydroxy-ester **4** (4611 mg, 90%) as a colorless oil. The product was characterized by ^1^H NMR and the data were found to be in good agreement with those published previously (84). The enantiomeric purity of compound **4** was determined as 99% *ee* through the synthesis of Mosher’s ester **11**, as described by Bauer *et al.* (29).

**(*R*)-3-Hydroxydecanoic acid (5)**

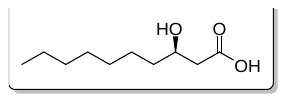

Alcohol **4** (4309 mg, 21.30 mmol, 1.00 eq) was solubilized in 1 M aq. NaOH (43 mL) at 0 °C. The mixture was stirred at 0 °C for 1 h, then at rt for an additional 1.5 h. The solution was acidified with 2 M aq. HCl until a pH ~2-3 was reached. The mixture was diluted in EtOAc and the aqueous phase was extracted with EtOAc (3×). The combined organic layers were washed with brine, dried over anhydrous MgSO_4_, filtered, and evaporated under reduced pressure to give acid **5** (3840 mg, 96%) as a white amorphous solid without further purification. The product was characterized by ^1^H NMR and the data were found to be in good agreement with those published previously (84). Purity of the compound was further confirmed by HPLC-CAD analysis using the previously described general method (9.0 minutes, see **supplementary chromatogram 1**).

**(*R*)-3-(*tert*-Butyldimethylsilyloxy)decanoic acid (6)**

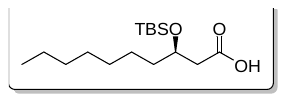

Imidazole (904 mg, 13.28 mmol, 10.0 eq) was added to a solution of TBSCl (701 mg, 4.65 mmol, 3.5 eq) in anhydrous DMF (2.3 mL) at 0 °C. The mixture was stirred at 0 °C for 15 min under an Ar atmosphere, after which a solution of acid **5** (252 mg, 1.33 mmol, 1.0 eq) in anhydrous DMF (0.5 mL) was added. The solution was stirred at rt for 16 h under an Ar atmosphere, then transferred to a separatory funnel. Brine was added to the mixture and the latter was extracted with a 1:3 mixture of Et_2_O/*n-*Hex. The organic layer was dried over anhydrous MgSO_4_, filtered, and evaporated under reduced pressure. The residue was solubilized in a mixture of MeOH (36 mL) and THF (18 mL) to which a solution of K_2_CO_3_ (450 mg, 3.26 mmol, 2.45 eq) in H_2_O (6 mL) was added at 0 °C. The mixture was stirred at 0 °C for 1 h after which brine (18 mL) was added. The solution was acidified with 1 M aq. HCl until a pH of 3 was reached. The solution was extracted with a 1:3 mixture of Et_2_O/*n-*Hex, dried over anhydrous MgSO_4_, filtered, and evaporated under reduced pressure. The residue was dried under high vacuum for 16 h to give acid **6** (402 mg, quant.) as a colorless oil. The product was characterized by ^1^H NMR and the data were found to be in good agreement with those published previously (84).

**(*R*)-Benzyl 3-hydroxydecanoate (7)**

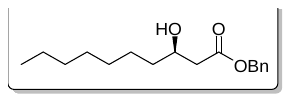

Acid **5** (84) (143 mg, 0.761 mmol, 1.0 eq) was solubilized in anhydrous DMF (3.8 mL) under Ar. BnBr (0.20 mL, 1.7 mmol, 2.2 eq) and Cs_2_CO_3_ (347 mg, 1.07 mmol, 1.4 eq) were successively added and the mixture was stirred for 21 h at rt under Ar. The suspension was diluted with DCM and washed with saturated aq. NH_4_Cl. The aqueous layer was extracted with DCM (2×). The combined organic layers were washed with brine, dried over anhydrous MgSO_4_, filtrated, and evaporated under reduced pressure. The residue was purified by silica gel flash chromatography (*n-*Hex/EtOAc 95:5 to 85:5) to give benzyl ester **7** (201 mg, quant.) as a colorless oil: *R_f_* 0.24 (*n-*Hex/EtOAc 8:2); [α]^20^_D_ –15 (*c* 0.6, CHCl_3_); ^1^H NMR (600 MHz, CDCl_3_) *δ* (ppm) 7.38−7.32 (m, 5H, 5 × C*H*_Bn_), 5.15 (s, 2H, C*H*_2Bn_), 4.04−4.01 (m, 1H, H-3), 2.85 (d, *J* = 3.5 Hz, 1H, O*H*), 2.56 (dd, *J_2a-2b_* = 16.5 Hz, *J*_2a-_*_3_* = 3.0 Hz, 1H, H-2a), 2.46 (dd, *J_2b-2a_* = 16.5 Hz, *J_2b-3_* = 9.1 Hz, 1H, H-2b), 1.54−1.26 (m, 12H, H-4, H-5, H-6, H-7, H-8, H-9), 0.88 (t, *J* = 7.0 Hz, 3H, H-10); ^13^C NMR (150 MHz, CDCl_3_) *δ* (ppm) 173.0 (C-1), 135.7 (C_Bn_), 128.8 (2C, 2 × *C*H_Bn_), 128.5 (*C*H_Bn_), 128.4 (2C, 2 x *C*H_Bn_), 68.2 (C-3), 66.6 (*C*H_2Bn_), 41.5 (C-2), 36.7−22.8 (6C, C-4, C-5, C-6, C-7, C-8, C-9), 14.2 (C-10); HRMS (ESI-TOF) *m/z* [M + Na]^+^ calcd for C_17_H_26_NaO_3_ 301.17742; found 301.17636.

**(*R*)-Benzyl 3-(((*R*)-3-((*tert*-butyldimethylsilyl)oxy)decanoyl)oxy)decanoate (8)**

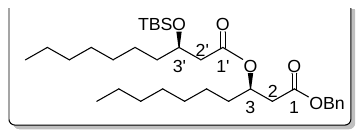

Benzyl ester **7** (91 mg, 0.32 mmol, 1.0 eq) and acid **6** (121 mg, 0.388 mmol, 1.2 eq) were solubilized in anhydrous DCE (3.9 mL) under Ar. EDC (186 mg, 0.970 mmol, 3.0 eq) and DMAP (12 mg, 0.10 mmol, 0.3 eq) were successively added and the mixture was stirred at rt for 16 h under Ar. The solution was evaporated under reduced pressure and the residue was purified by silica gel flash chromatography (*n-*Hex/EtOAc 95:5) to give dilipid **8** (171 mg, 93%) as a colorless oil: *R_f_* 0.57 (*n-*Hex/EtOAc 8:2); [α]^20^_D_ +53 (*c* 0.7, CHCl_3_); ^1^H NMR (600 MHz, CDCl_3_) *δ* (ppm) 7.37−7.31 (m, 5H, 5 × C*H*_Bn_), 5.24−5.20 (m, 1H, H-3), 5.11 (s, 2H, C*H*_2Bn_), 4.07 (p, *J* = 6.8 Hz, 1H, H-3’), 2.65 (dd, *J_2a-2b_* = 15.4 Hz, *J_2a-3_* = 7.1 Hz, 1H, H-2a), 2.57 (dd, *J_2b-2a_* = 15.4 Hz, *J_2b-3_* = 5.8 Hz, 1H, H-2b), 2.41 (dd, *J_2a’-2b’_* = 14.8 Hz, *J_2a’-3’_* = 5.9 Hz, 1H, H-2a’), 2.36 (dd, *J_2b’-2a’_* = 14.8 Hz, *J_2b’-3’_* = 6.7 Hz, 1H, H-2b’), 1.63−1.24 (m, 24H, 12 × C*H*_2_), 0.89−0.86 (m, 15H, H-10, H-10’, C(C*H*_3_)_3TBS_), 0.06 (s, 3H, C*H*_3TBS_), 0.04 (s, 3H, C*H*_3TBS_); ^13^C NMR (150 MHz, CDCl_3_) *δ* (ppm) 171.1, 170.3 (2C, C-1, C-1’), 135.9 (C_Bn_), 128.7, 128.4 (5C, 5 × *C*H_Bn_), 70.7 (C-3), 69.4 (C-3’), 66.6 (C*H*_2Bn_), 42.9 (C-2’), 39.3 (C-2), 37.5−29.3 (8C, 8 × *C*H_2_), 26.0 (3C, C(*C*H_3_)_3TBS_), 25.3−22.8 (4C, 4 × *C*H_2_), 18.2 (*C*(CH_3_)_3TBS_), 14.3, 14.2 (2C, C-10, C-10’), −4.46 (2C, 2 × *C*H_3TBS_); HRMS (ESI-TOF) *m/z* [M + Na]^+^ calcd for C_17_H_26_NaO_3_ 585.39457; found 585.39279.

**(*R*)-Benzyl 3-(((*R*)-3-hydroxydecanoyl)oxy)decanoate (9)**

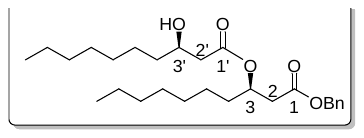

A solution of dilipid **8** (192 mg, 0.340 mmol, 1.0 eq) in DCM (0.7 mL) was added dropwise in TFA (1.4 mL) during a one-minute period, then the mixture was stirred for one additional minute. The reaction mixture was quenched with saturated aq. NaHCO_3_. The organic layer was dried over anhydrous MgSO_4_, filtered, and evaporated under reduced pressure. The residue was purified by silica gel flash chromatography (*n-*Hex/EtOAc 9:1 to 8:2) to give alcohol **9** (143 mg, 94%) as a colorless oil: *R_f_* 0.28 (*n-*Hex/EtOAc 8:2); [α]^20^_D_ –13 (*c* 0.4, CHCl_3_); ^1^H NMR (600 MHz, CDCl_3_) *δ* (ppm) 7.38−7.32 (m, 5H, 5 × C*H*_Bn_), 5.30-5.26 (m, 1H, H-3), 5.11 (s, 2H, C*H*_2Bn_), 3.98−3.94 (m, 1H, H-3’), 2.95 (br s, 1H, OH), 2.63 (dd, *J_2a-2b_* = 14.2 Hz, *J_2a-3_* = 6.2 Hz, 1H, H-2a), 2.60 (dd, *J_2b-2a_* = 14.2 Hz, *J_2b-3_* = 4.1 Hz, 1H, H-2b), 2.42 (dd, *J_2a’-2b’_* = 15.8 Hz, *J_2a’-3’_* = 2.9 Hz, 1H, H-2a’), 2.31 (dd, *J_2b’-2a’_* = 15.8 Hz, *J_2b’-3’_* = 9.2 Hz, 1H, H-2b’), 1.64−1.25 (m, 24H, 12 × C*H*_2_), 0.89−0.85 (m, 6H, H-10, H-10’); ^13^C NMR (150 MHz, CDCl_3_) *δ* (ppm) 172.6, 170.6 (2C, C-1, C-1’), 135.7 (C_Bn_), 128.7 (2C, 2 × *C*H_Bn_), 128.6 (2C, 2 × *C*H_Bn_), 128.5 (*C*H_Bn_), 71.0 (C-3), 68.4 (C-3’), 66.8 (*C*H_2Bn_), 41.9 (C-2’), 39.3 (C-2), 36.7−22.8 (12C, 12 × *C*H_2_), 14.2 (2C, C-10, C-10’); HRMS (ESI-TOF) *m/z* [M + Na]^+^ calcd for C_27_H_44_NaO_5_ 471.3081; found 471.3089; *m/z* [M + K]^+^ calcd for C_27_H_44_O_5_K 487.28203; found 487.28242.

**HAA (C_10_-C_10_) (10)**

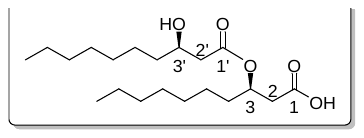

Alcohol **9** (29 mg, 0.064 mmol, 1.0 eq) was solubilized in MeOH (1.5 mL) under an Ar atmosphere. Pd black (29 mg, 1 mg⋅mg^–1^ of alcohol **9**) was added and the suspension was stirred at 40 °C under H_2_ atmosphere for 16 h. The suspension was filtered over Celite and evaporated under reduced pressure. The residue was purified by silica gel flash chromatography (DCM/MeOH 95:5) to give dilipid **10** (20 mg, 90%) as a colorless oil: *R_f_* 0.22 (DCM/MeOH 95:5); [α]^20^_D_ –18 (*c* 0.3, CHCl_3_); ^1^H NMR (600 MHz, CDCl_3_) *δ* (ppm) 5.29 (m, 1H, H-3), 4.03 (br s, 1H, H-3’), 2.60 (br s, 2H, H-2), 2.48 (dd, *J_2a’-2b’_* = 15.7 Hz, *J_2a’-3’_* = 2.3 Hz, 1H, H-2a’), 2.40 (dd, *J_2b’-2a’_* = 15.8 Hz, *J_2b’-3’_* = 9.3 Hz, 1H, H-2b’), 1.64−1.26 (m, 24H, 12 × C*H*_2_), 0.88 (distorted t, *J* = 7.0 Hz, 6H, H-10, H-10’); ^13^C NMR (150 MHz, CDCl_3_) *δ* (ppm) 172.7 (2C, 2 x *C*OOR), 71.0 (C-3), 68.6 (C-3’), 41.9 (2C, C-2, C-2’), 36.7−22.8 (12C, 12 × *C*H_2_), 14.24, 14.22 (2C, C-10, C-10’). Purity of the compound was further confirmed by HPLC-CAD analysis using the previously described general method (42.8 minutes, see **supplementary chromatogram 1**).

**Determination of the enantiomeric purity of methyl ester 4 through its derivatization as (*R*)-methyl α-methoxy-α-trifluoromethylphenylacetyl-(*R*)-3-hydroxydecanoate (11)**

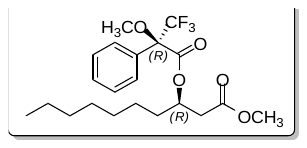

Methyl ester **3** (84) (15 mg, 0.074 mmol, 1.0 eq) was solubilized in anhydrous DCE (0.4 mL). Mosher’s acid (26 mg, 0.11 mmol, 1.5 eq), EDC (28 mg, 0.15 mmol, 2.0 eq) and DMAP (2 mg, 0.02 mmol, 0.2 eq) were successively added to the solution, which was then stirred at rt for 16 h under an Ar atmosphere. The solution was evaporated under reduced pressure and the residue was purified by silica gel flash chromatography (DCM) to give Mosher’s ester **11 (**29) (15 mg, 50%) as a colorless oil. The product was characterized by ^1^H NMR and the data were found to be in good agreement with those published previously. Analysis of this spectra allowed the determination of the enantiomeric purity of methyl ester **3** (99% e.r.), as reported by Bauer *et al.* (29).

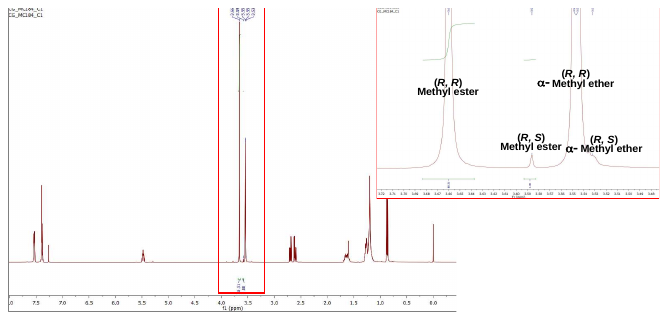

Enlargement of the ^1^H NMR spectrum of Mosher’s ester **11**.
