## Supplementary data 2 for "The bacterial virulence factors rhamnolipids and their (*R*)-3-hydroxyalkanoate precursors activate *Arabidopsis* innate immunity through two independent mechanisms"

### Spectra of novel synthetic compounds

### Supplementary spectrum 1 | ^1^H NMR spectrum (CDCl_3_, 600 MHz) of (*R*)-benzyl 3-hydroxydecanoate (7)

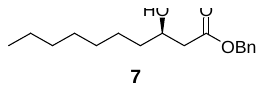

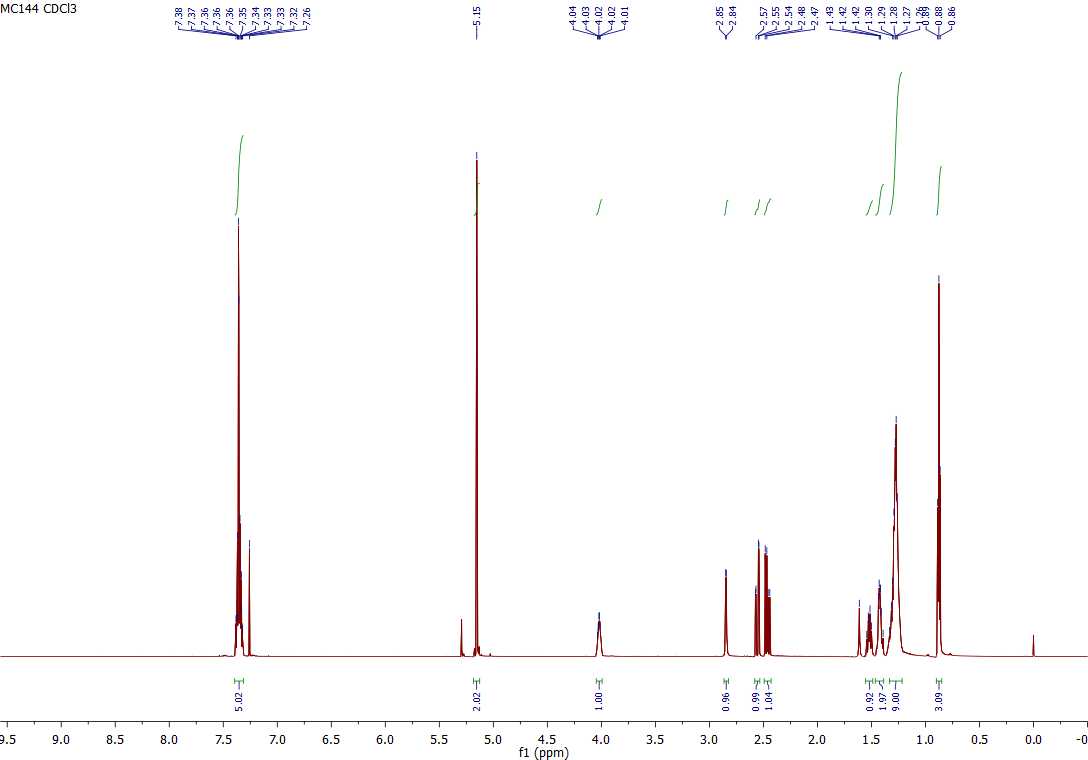

### Supplementary spectrum 2 | COSY NMR spectrum (CDCl_3_, 600 MHz) of (*R*)-benzyl 3-hydroxydecanoate (7)

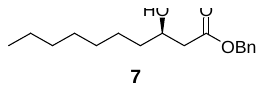

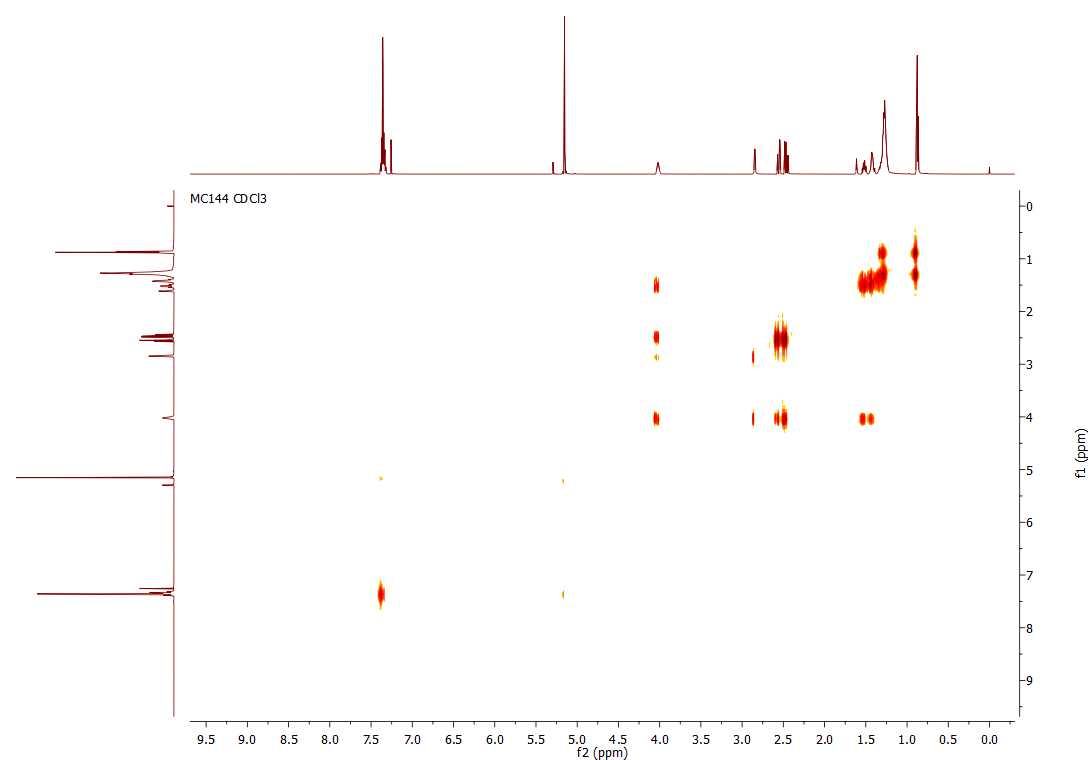

### Supplementary spectrum 3 | ^13^C NMR spectrum (CDCl_3_, 150 MHz) of (*R*)-benzyl 3-hydroxydecanoate (7)

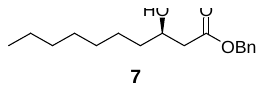

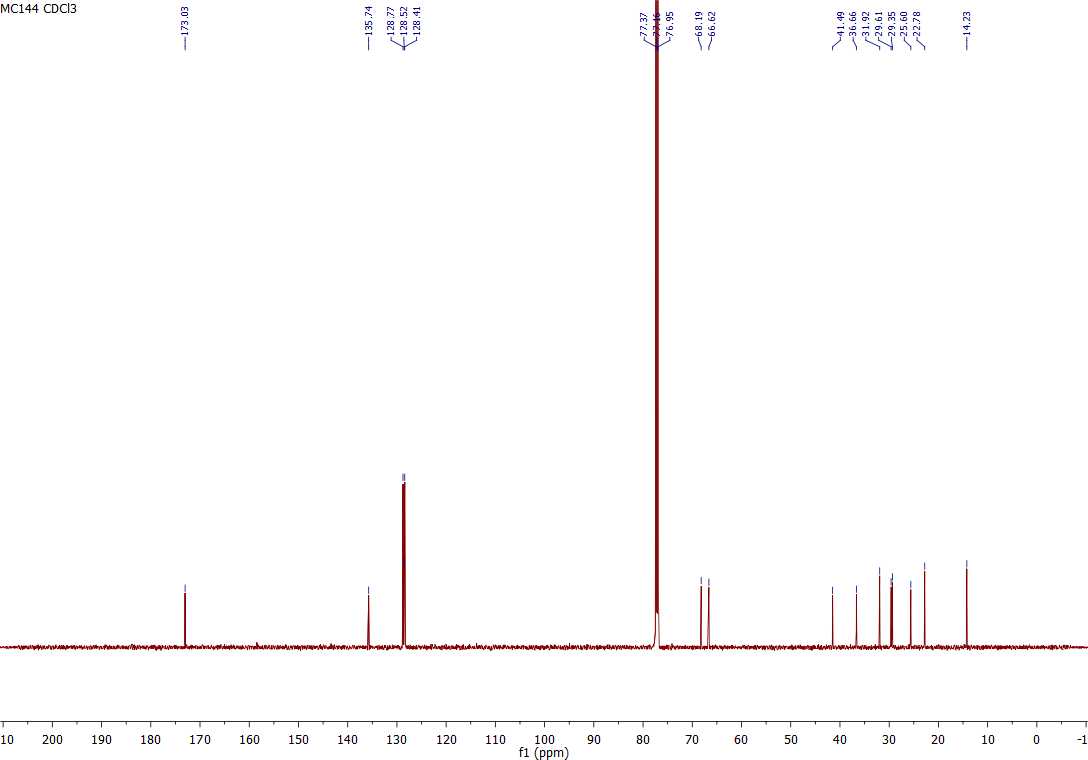

### Supplementary spectrum 4 | ^­^HSQC NMR spectrum (CDCl_3_, 600 MHz) of (*R*)-benzyl 3-hydroxydecanoate (7)

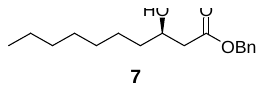

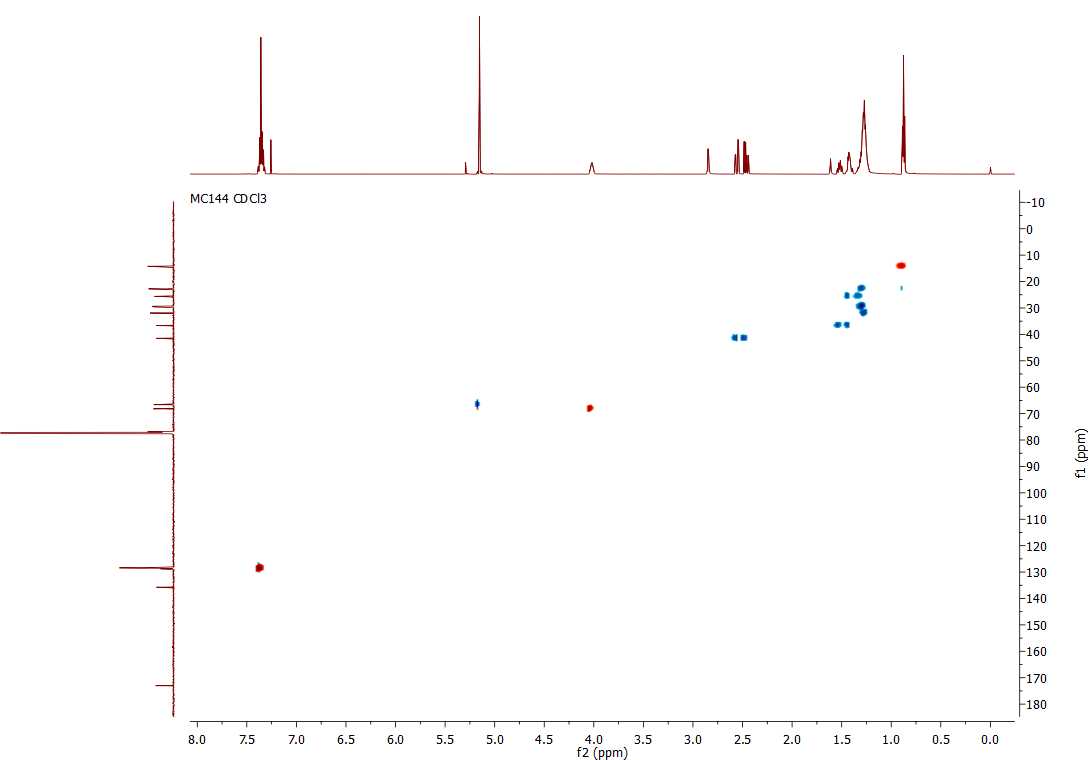

### Supplementary spectrum 5 | ^­1^H NMR spectrum (CDCl_3_, 600 MHz) of (*R*)-benzyl 3-(((*R*)-((*tert*-butyldimethylsilyl)oxy)decanoyl)oxy)decanoate (8)

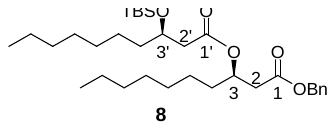

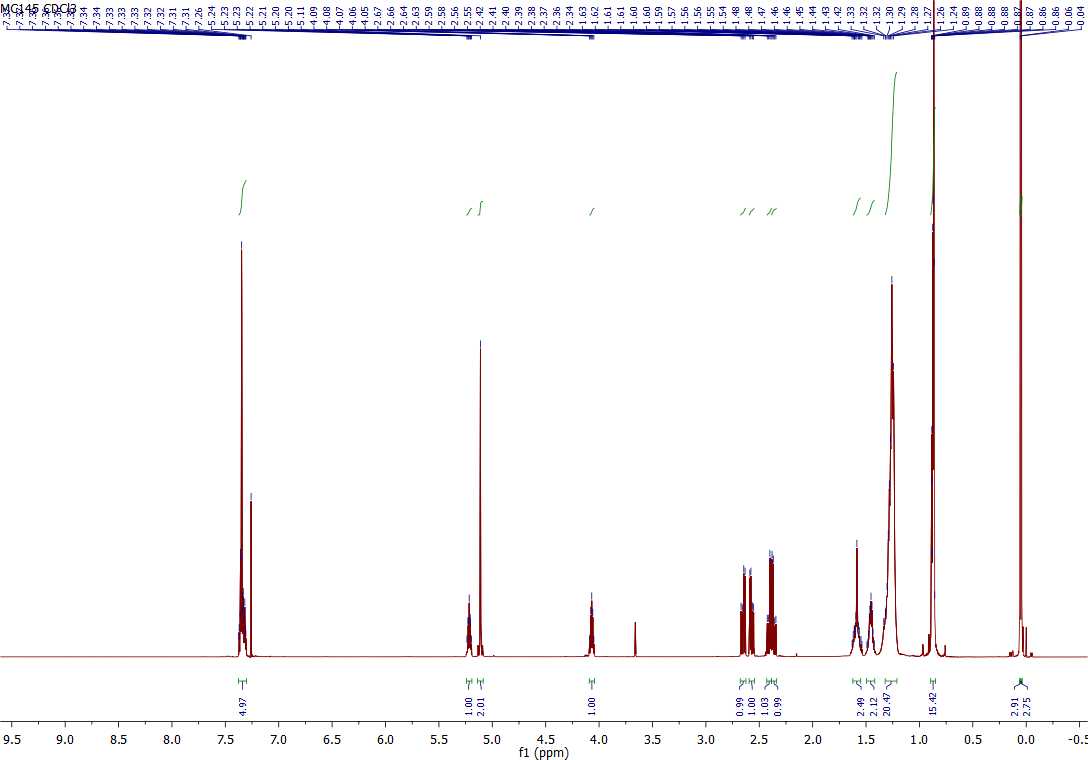

### Supplementary spectrum 6 | COSY NMR spectrum (CDCl_3_, 600 MHz) of (*R*)-benzyl 3-(((*R*)-((*tert*-butyldimethylsilyl)oxy)decanoyl)oxy)decanoate (8)

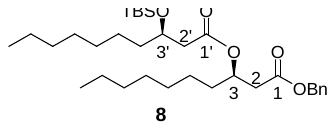

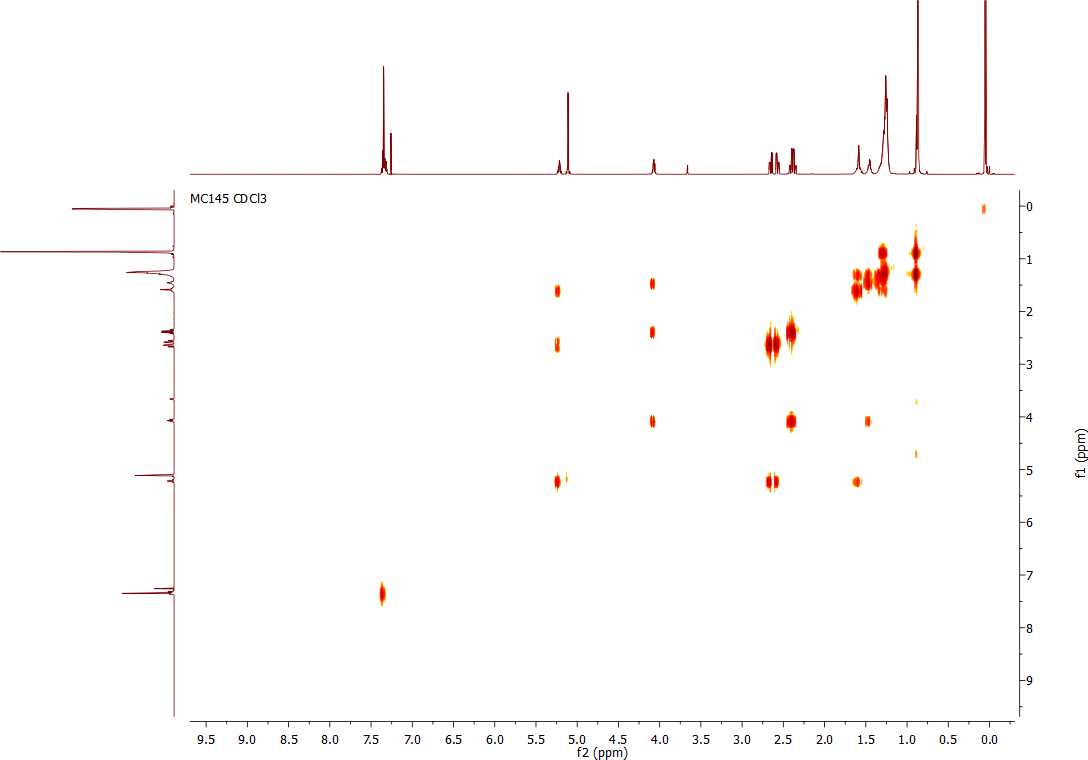

### Supplementary spectrum 7 | ^­13^C NMR spectrum (CDCl_3_, 150 MHz) of (*R*)-benzyl 3-(((*R*)-((*tert*-butyldimethylsilyl)oxy)decanoyl)oxy)decanoate (8)

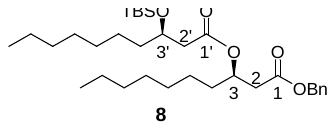

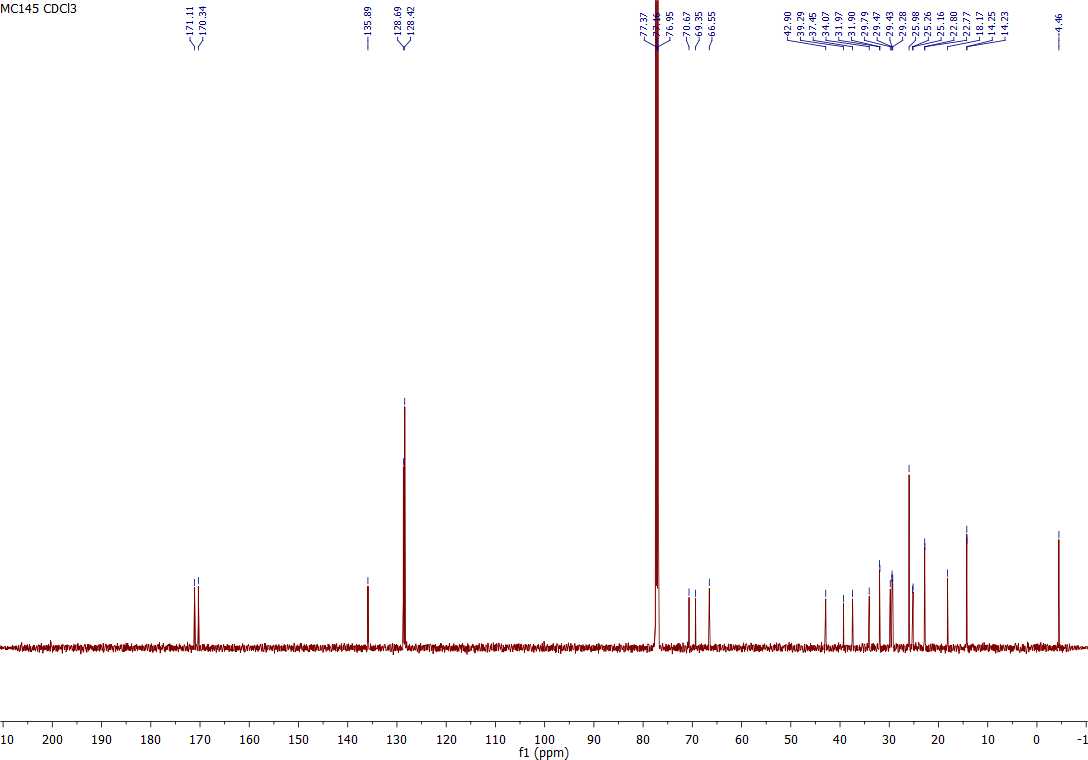

### Supplementary spectrum 8 | ^­^HSQC NMR spectrum (CDCl_3_, 600 MHz) of (*R*)-benzyl 3-(((*R*)-((*tert*-butyldimethylsilyl)oxy)decanoyl)oxy)decanoate (8)

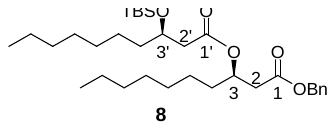

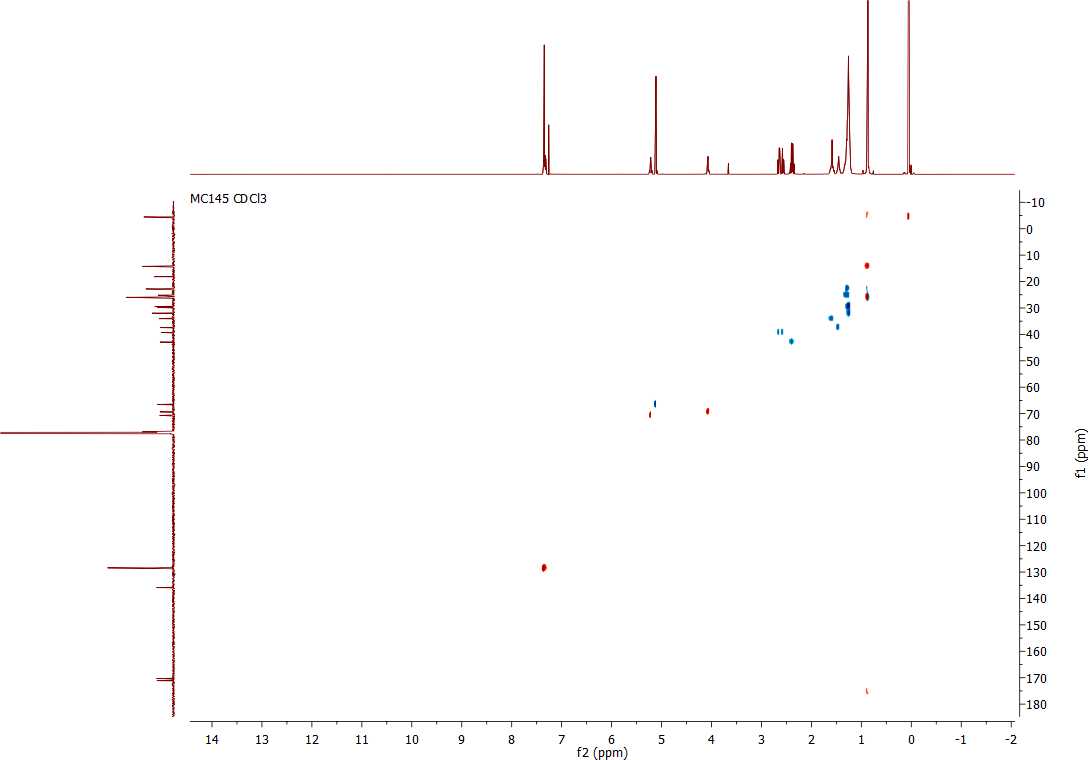

### Supplementary spectrum 9 | ^­1^H NMR spectrum (CDCl_3_, 600 MHz) of (*R*)-benzyl 3-(((*R*)-3-hydroxydecanoyloxy)decanoate (9)

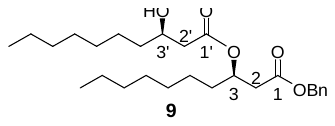

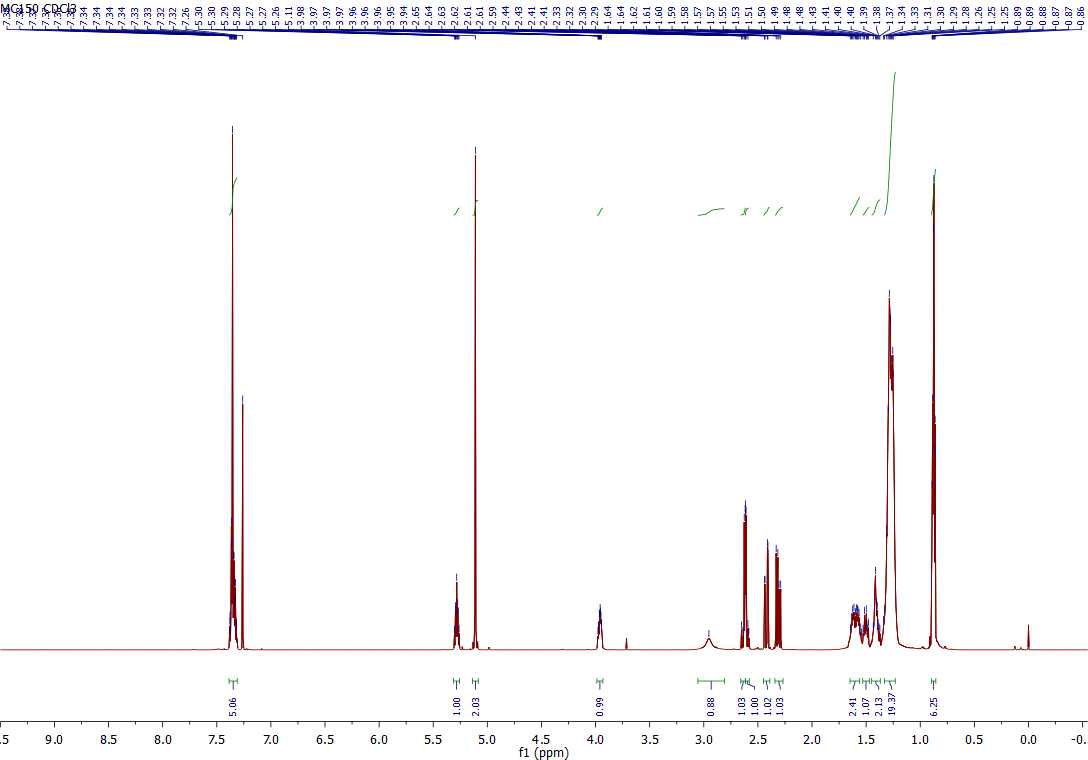

### Supplementary spectrum 10 | COSY NMR spectrum (CDCl_3_, 600 MHz) of (*R*)-benzyl 3-(((*R*)-3-hydroxydecanoyloxy)decanoate (9)

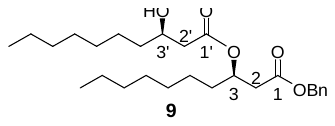

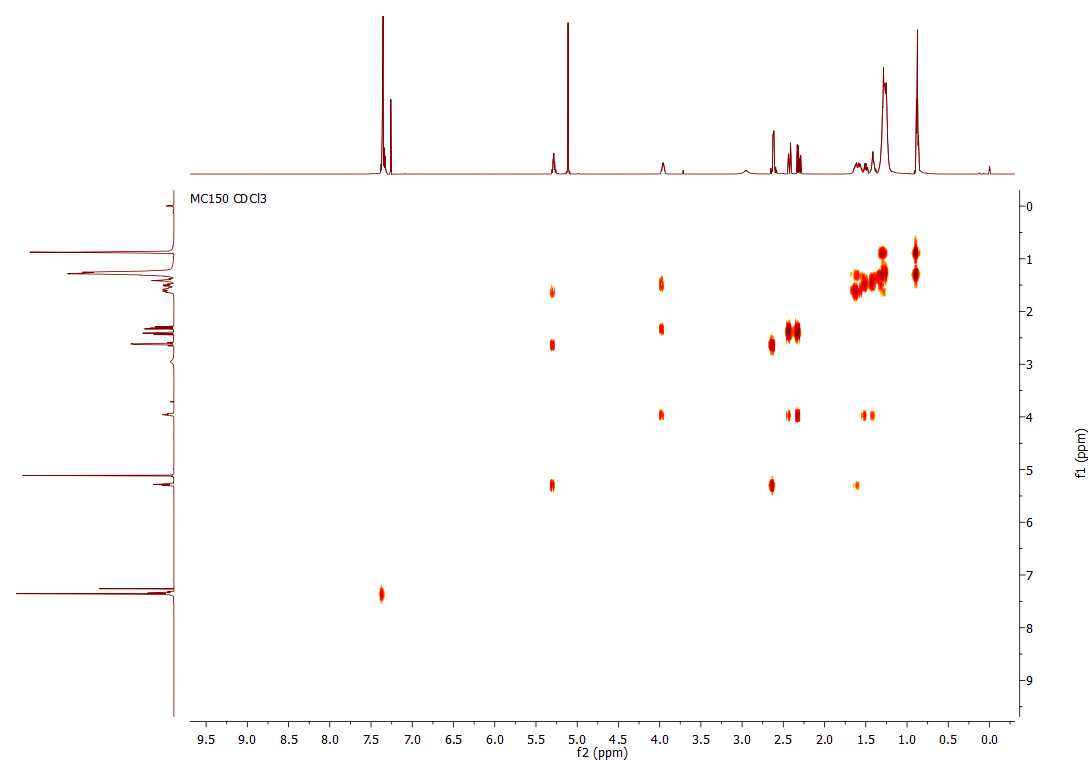

### Supplementary spectrum 11 | ^­13^C NMR spectrum (CDCl_3_, 150 MHz) of (*R*)-benzyl 3-(((*R*)-3-hydroxydecanoyloxy)decanoate (9)

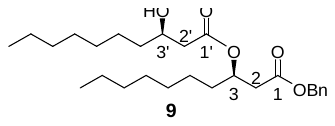

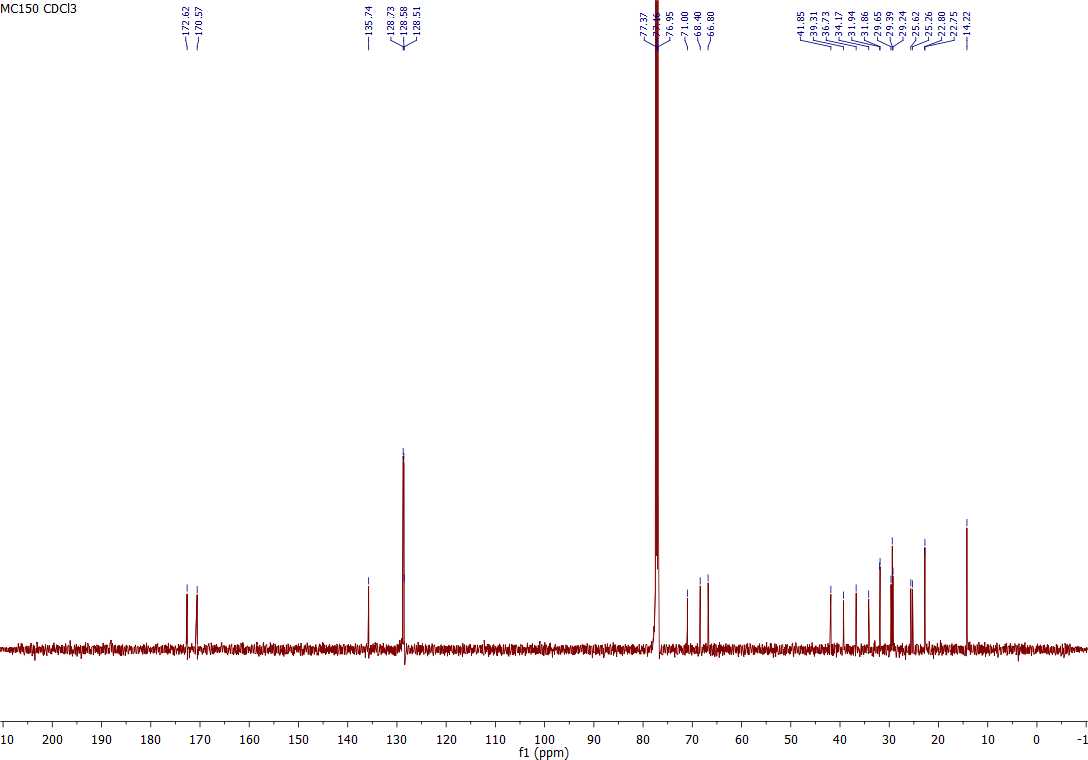

### Supplementary spectrum 12 | HSQC NMR spectrum (CDCl_3_, 600 MHz) of (*R*)-benzyl 3-(((*R*)-3-hydroxydecanoyloxy)decanoate (9)

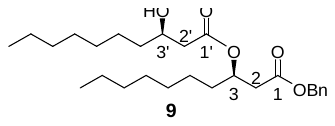

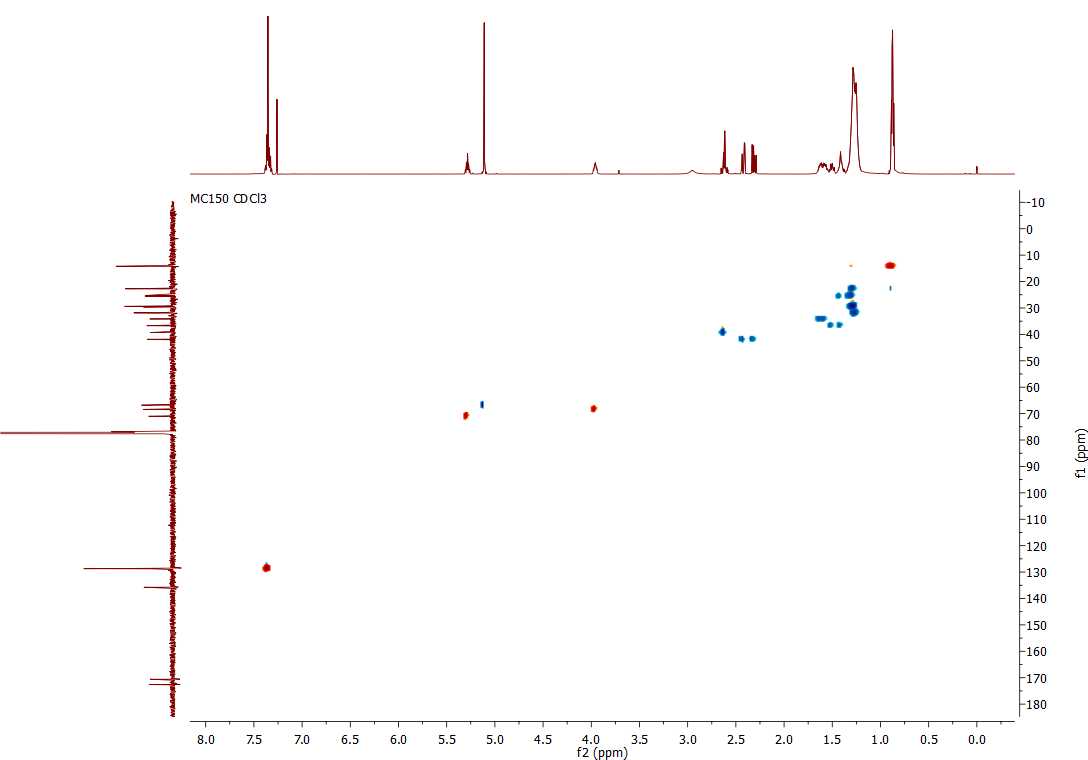

### Supplementary spectrum 13 | ^1^H NMR spectrum (CDCl_3_, 600 MHz) of 3-(3-hydroxydecanoyloxy)decanoic acid (10)

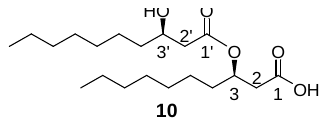

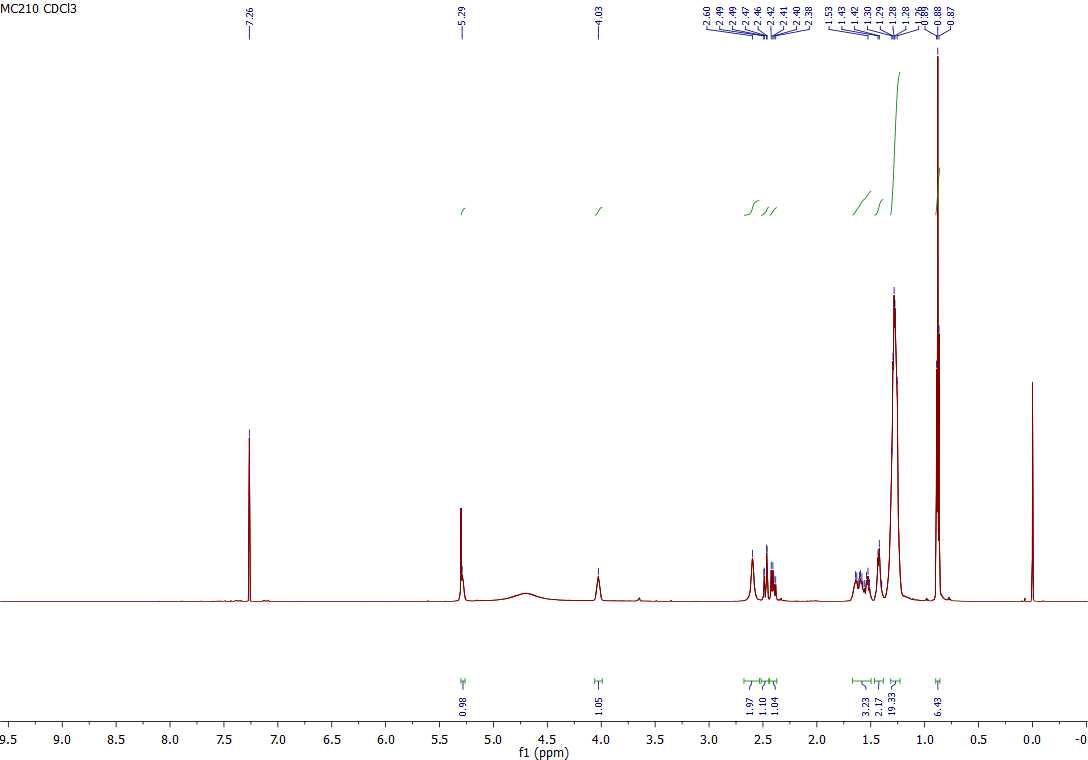

### Supplementary spectrum 14 | COSY NMR spectrum (CDCl_3_, 600 MHz) of 3-(3-hydroxydecanoyloxy)decanoic acid (10)

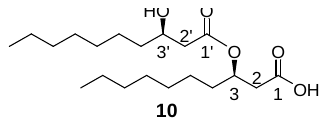

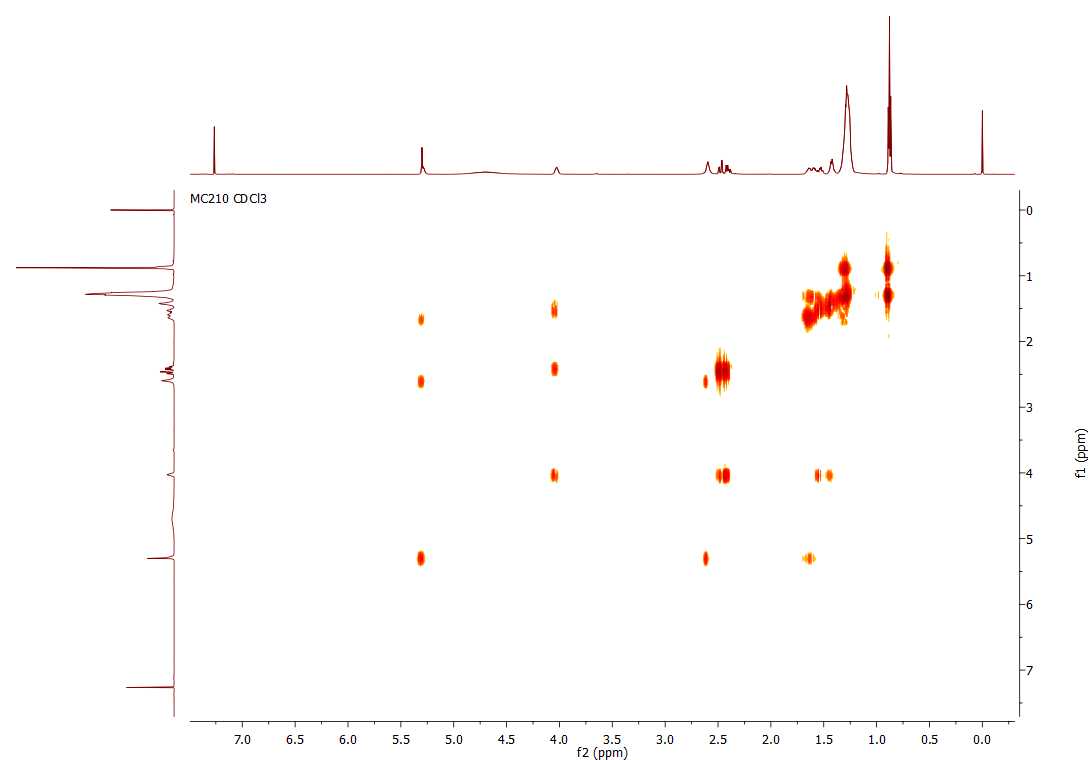

### Supplementary spectrum 15 | ^13^C NMR spectrum (CDCl_3_, 150 MHz) of 3-(3-hydroxydecanoyloxy)decanoic acid (10)

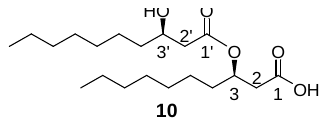

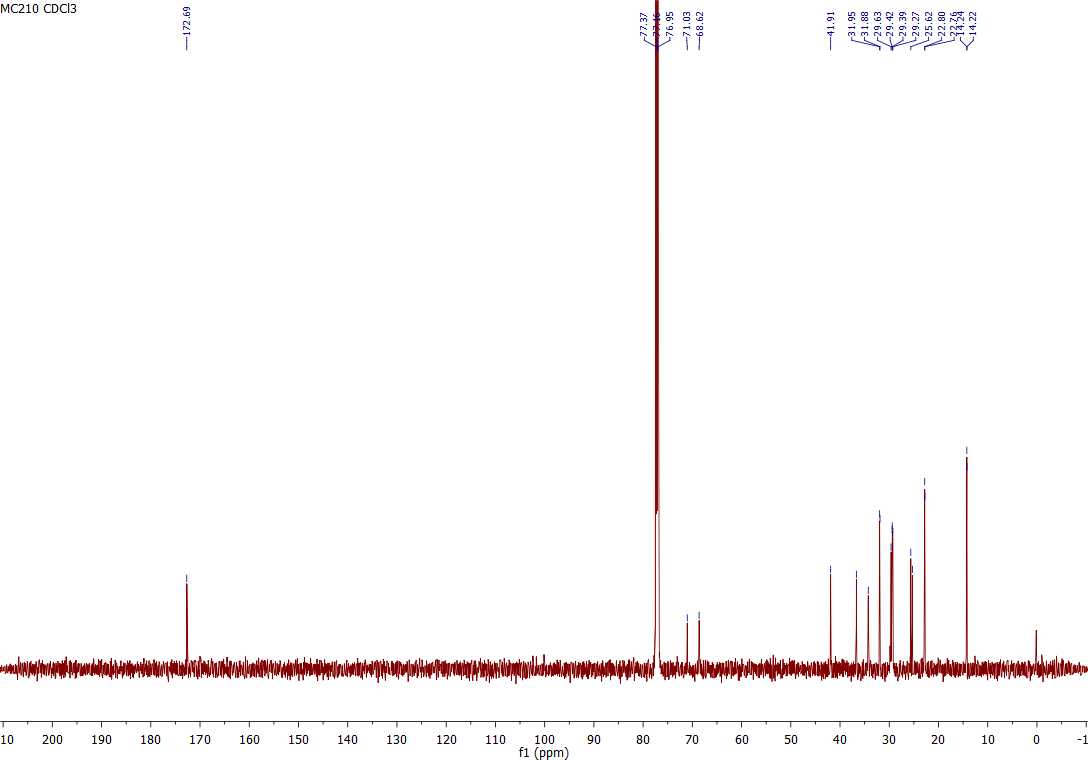

### Supplementary spectrum 16 | HSQC NMR spectrum (CDCl_3_, 600 MHz) of 3-(3-hydroxydecanoyloxy)decanoic acid (10)

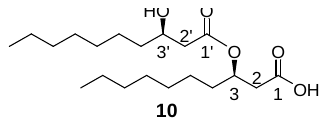

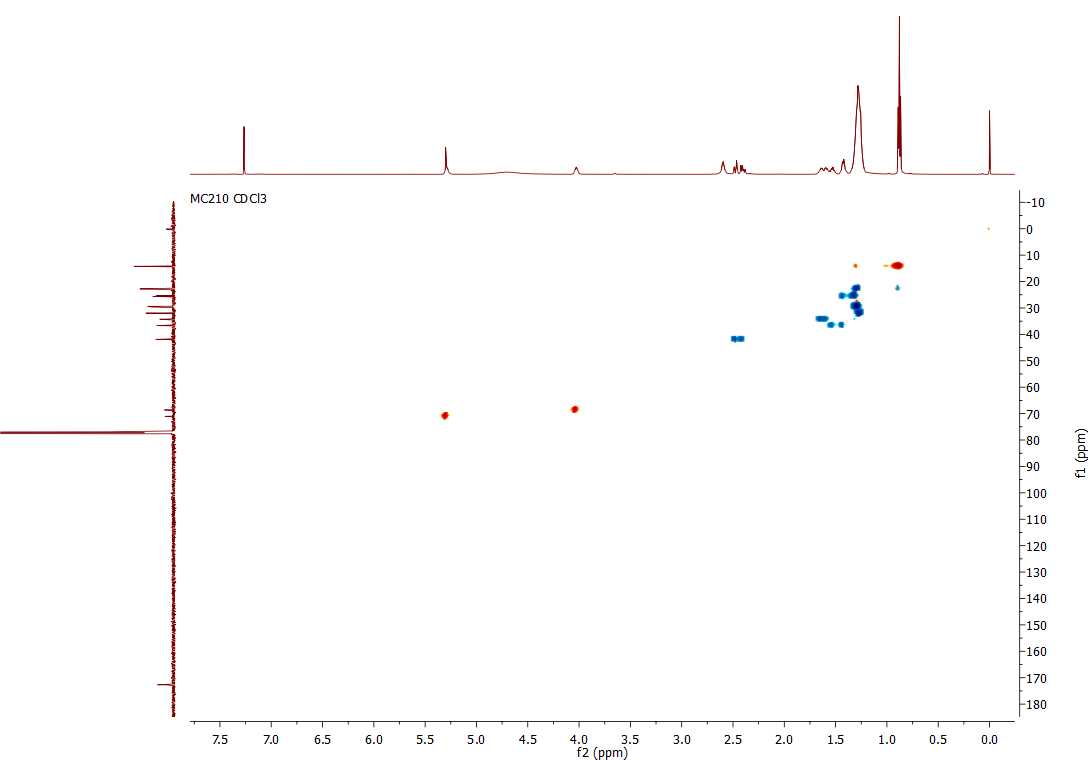

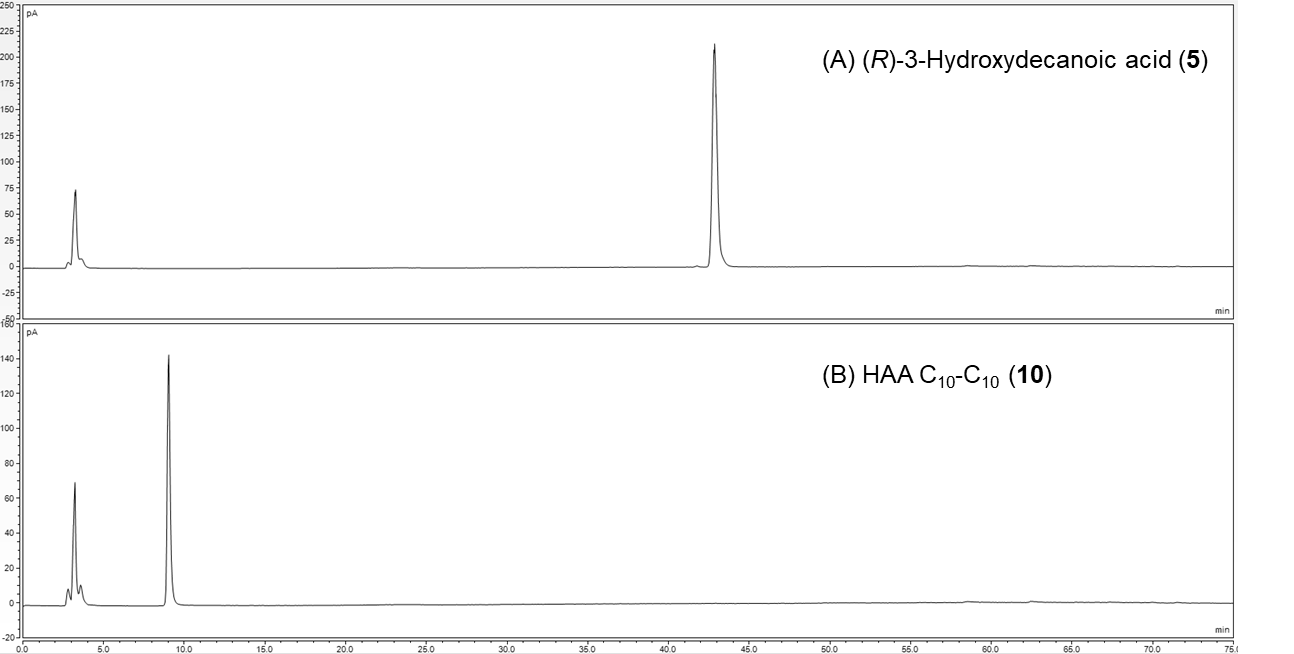

**Supplementary chromatogram 1.**  HPLC-CAD chromatograms of (A) synthetic (*R*)-3-hydroxydecanoic acid **5** and (B) synthetic HAA C_10_-C_10_ **10**
